# MemBack: An Equivariant Graph Neural Network for Backmapping Lipid Membranes

**DOI:** 10.64898/2026.08.19.745274

**Authors:** Y. Eren Tunç, Rainer Böckmann

## Abstract

Backmapping coarse-grained simulations to atomistic resolution is central to multiscale molecular simulation but remains challenging for chemically complex lipid membranes. We introduce *MemBack*, an SE(3)-equivariant graph neural network that reconstructs CHARMM36 lipid structures from Martini 3 configurations by single-pass heavy-atom prediction followed by automated post-processing. Across chemically diverse systems, *MemBack* achieved a mean superposition-free per-lipid heavy-atom RMSD of 0.65 Å, while retaining comparable accuracy for membrane systems and lipid species excluded from training. At the membrane scale, *MemBack* preserved structural organization across resolutions: in a 16-component red-blood-cell membrane model excluded from training, bilayer thickness, area per lipid, and acyl-chain order closely matched the atomistic reference after only restrained minimization; in a native phase-separated Martini 3 membrane, the lateral organization of ordered and disordered domains was likewise retained directly after backmapping, without atomistic equilibration. Native Martini 3 systems containing up to 1.4 million reconstructed atoms remained consistent with their parent coarse-grained configurations and could be propagated in atomistic simulations, providing an efficient interface between Martini 3 and CHARMM36 membrane simulations.

## 1 Introduction

Molecular dynamics (MD) simulations have become an indispensable tool for obtaining molecular-level insight into the structure, dynamics, and thermodynamics of biomembranes [1]. This level of structural and dynamical detail is rarely accessible to experiments alone. By numerically solving Newton’s equations of motion for each particle in the system, MD provides a time-resolved, atomistic view of processes such as lipid self-assembly, membrane phase transitions, and protein-lipid interactions. The principal drawback of the approach is its limited accessible scale: chemically detailed, all-atom (AA) force fields require integration time steps on the femtosecond scale, while many biologically relevant processes (domain formation, large-scale membrane remodeling, and lateral lipid reorganization) occur over microsecond to millisecond timescales and extend across tens to hundreds of nanometers. Direct atomistic sampling of these regimes remains prohibitively costly, even with specialized computational hardware.

Coarse-grained (CG) models address this limitation by reducing the number of degrees of freedom in the system. In the widely used Martini framework [2, 3], groups of typically three to six heavy atoms are mapped onto a single interaction site, or bead. The resulting reduction in particle number, together with a simplified treatment of electrostatics, the increased mass of the CG beads, and the smoother effective energy landscape arising from the averaging of atomistic interactions, substantially lowers the computational cost. In particular, the increased bead masses and the absence of fast atomistic degrees of freedom permit integration time steps of approximately 20 fs, about an order of magnitude larger than those commonly used in all-atom (AA) simulations. Combined, these features yield effective speed-ups of two to three orders of magnitude relative to AA simulations, enabling access to substantially larger spatial and temporal scales. This gain comes at the expense of chemical resolution: properties that depend explicitly on atomic coordinates, including hydrogen-bond geometries, detailed dihedral-angle distributions, atomistically resolved order parameters, or quantities requiring an electronic-structure description, are no longer directly accessible from a CG trajectory. Atomistic and coarse-grained representations are therefore complementary rather than competing: AA models provide detailed chemical information over comparatively limited spatial and temporal scales, whereas CG models enable the investigation of larger-scale and longer-timescale dynamics at reduced resolution.

A natural strategy for combining these complementary strengths is a multiscale workflow in which slow, large-scale processes are first sampled at CG resolution, followed by reconstruction of selected configurations at full atomistic detail for validation, refinement, and further analysis (see, e.g. [4, 5]). The transformation from a coarse-grained to an atomistic representation is commonly referred to as backmapping. This inverse problem is intrinsically underdetermined: because multiple atoms are represented by a single CG bead, a given CG configuration does not specify a unique atomistic structure but is compatible with an ensemble of atomistic configurations. Backmapping therefore requires the reconstruction of a high-dimensional atomistic geometry from a strongly compressed and information-lossy representation, while simultaneously producing structures that are chemically plausible, free of severe steric clashes, and compatible with the target atomistic force field.

A range of strategies have been developed to address this problem, with geometric and fragment-based approaches currently prevailing [6–8]. The Backward [6] protocol carries out a smart initial placement of atoms using the coordinates of nearby beads, followed by successive force-field energy minimizations and restrained MD steps to resolve unfavorable interactions and relax the reconstructed structure. CG2AT2 [7], in contrast, reconstructs molecules using stereochemically defined atomistic fragments that are aligned with the corresponding CG representation before subsequent refinement. More recently, machine-learning approaches have been incorporated into fragment-based reconstruction. ART-SM [9], for example, learns atomistic conformational distributions from simulation data and selects fragment conformations that are compatible with the local CG geometry, followed by refinement of bonds, angles, and dihedrals. In parallel, fully data-driven generative approaches have emerged that reconstruct atomistic coordinates without relying on predefined fragment libraries, including conditional variational autoencoders [10], equivariant graph neural networks such as HEroBM [11], and flow-matching approaches such as FlowBack [12]. MSBack [13] extends generative backmapping to highly coarse-grained protein representations, using constrained diffusion to reconstruct atomistic detail consistent with the CG coordinates and combining this with physics-based reconstruction at finer resolution. Despite these advances, learned backmapping approaches are often tailored to particular molecular classes or CG representations, have been validated predominantly for proteins or small molecules, or require multiple computationally expensive refinement or sampling steps when generating atomistic structures. For complex lipid bilayers, comparatively little work has established whether learned backmapping can recover atomistically realistic local structure while simultaneously preserving membrane-level properties, including lipid density profiles, bilayer thickness, area per lipid, acyl-chain order, lateral packing, and the existence of ordered or disordered membrane domains, without requiring extensive subsequent atomistic relaxation.

In this work, we develop a graph neural network (GNN)-based approach for backmapping Martini 3 lipid systems to the CHARMM lipid force field [14–16]. Each CG configuration is represented as a graph in which beads constitute nodes carrying chemical information, while bonded interactions define edges with associated geometric features. We evaluate the model across chemically distinct membrane compositions and benchmark the reconstructed atomistic structures against reference simulations using local geometric and conformational metrics, together with membrane-level observables including bilayer thickness, area-per-lipid distributions, and domain formation. The resulting model provides a transferable, single-pass route from coarse-grained (Martini 3) lipid assemblies to atomistic representations (CHARMM36), recovering local atomistic structure while preserving the collective organization of complex lipid systems.

## 2 Methodology

MemBack reconstructs atomistic structures of lipid assemblies from Martini 3 coarse-grained (CG) structures in a single forward pass of a graph neural network (GNN). The predicted atomistic coordinates are subsequently processed using a deterministic pipeline that adds hydrogen atoms, enforces correct stereochemistry, resolves residual steric clashes, converts CG water and ion beads to their atomistic representations, and assembles the result into a simulation-ready system. The workflow comprises three stages. In the training stage, carried out once, atomistic reference structures are forward-mapped to Martini 3 resolution and converted into molecular graphs whose nodes are CG beads and whose edges follow the bonded topology. These graph pairs are used to train the equivariant network to predict the heavy-atom displacements belonging to each bead. In the prediction stage, applied to any new CG structure, the trained network predicts heavy-atom positions in a single forward pass, followed by the deterministic post-processing steps above. At the simulation stage, the system is energy minimized, equilibrated and used as a starting point for production simulation. The complete workflow is summarized in Figure 1.

**Figure 1:**
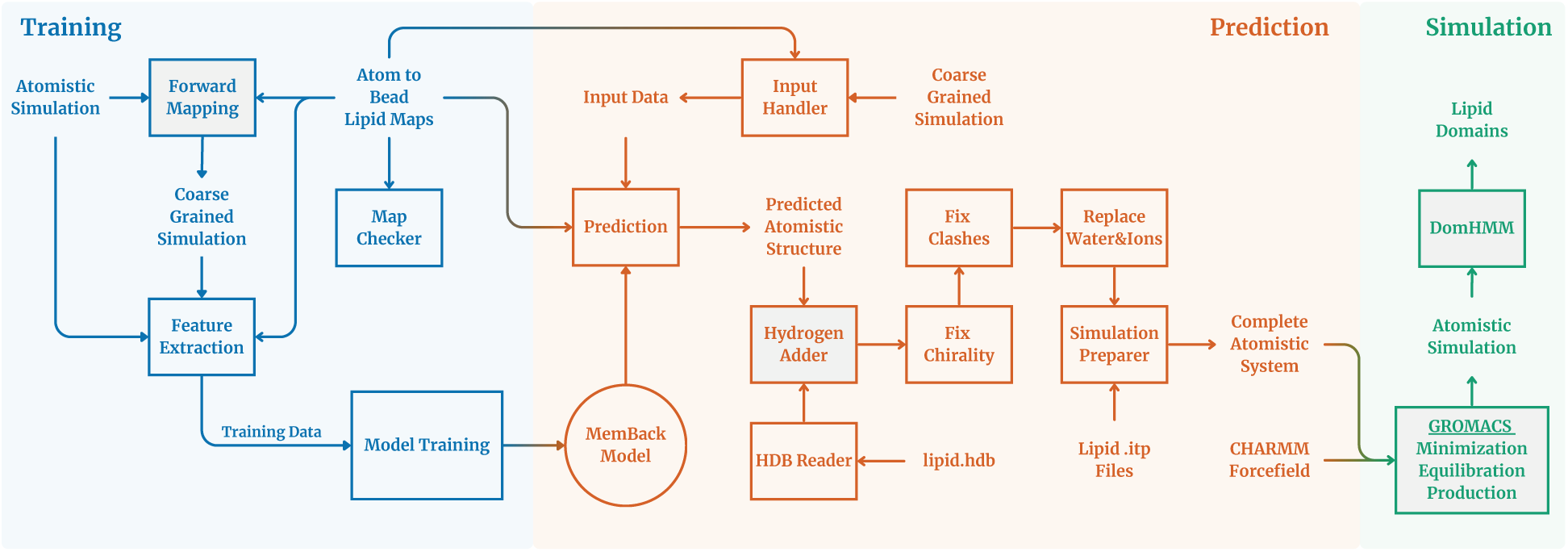
Overview of the MemBack workflow. Training (once, blue): all-atom (AA) reference structures are forward-mapped to coarse-grained (CG) resolution and converted into molecular graphs, which are used to train the equivariant GNN. Prediction (per system, orange): a CG structure is encoded as a graph, heavy-atom positions are predicted in a single forward pass, and the deterministic steps (hydrogen reconstruction, solvent and ion conversion, chirality correction, clash resolution and system assembly) yield the final atomistic system. Simulation (green): the assembled system is energy-minimized with restraints, equilibrated and used directly as the starting point for production atomistic molecular dynamics. Gray boxes denote steps involving third-party software.

### 2.1 Coarse-to-atomistic mapping

Reference all-atom (AA) CHARMM36 lipid trajectories were converted to their corresponding Martini 3 coarse-grained (CG) representations by forward mapping with PyCGTOOL [17], using bead-to-atom mapping files (.map). For each lipid residue type, the mapping file specifies the Martini bead names and types, bead charges, and the set of heavy atoms assigned to each bead. Mapping files were generated for the Martini 3 lipid species considered in this work.

Because both manually and automatically generated mapping files are susceptible to errors, each candidate mapping was subjected to an automated consistency check. The validation script compares the reference AA system with the CG system obtained by forward mapping and identifies duplicated atoms, atoms missing from the mapping, atom names absent from the AA topology, and bead names absent from the CG topology. Representative forward-mapped structures are shown in Figure 2.

**Figure 2:**
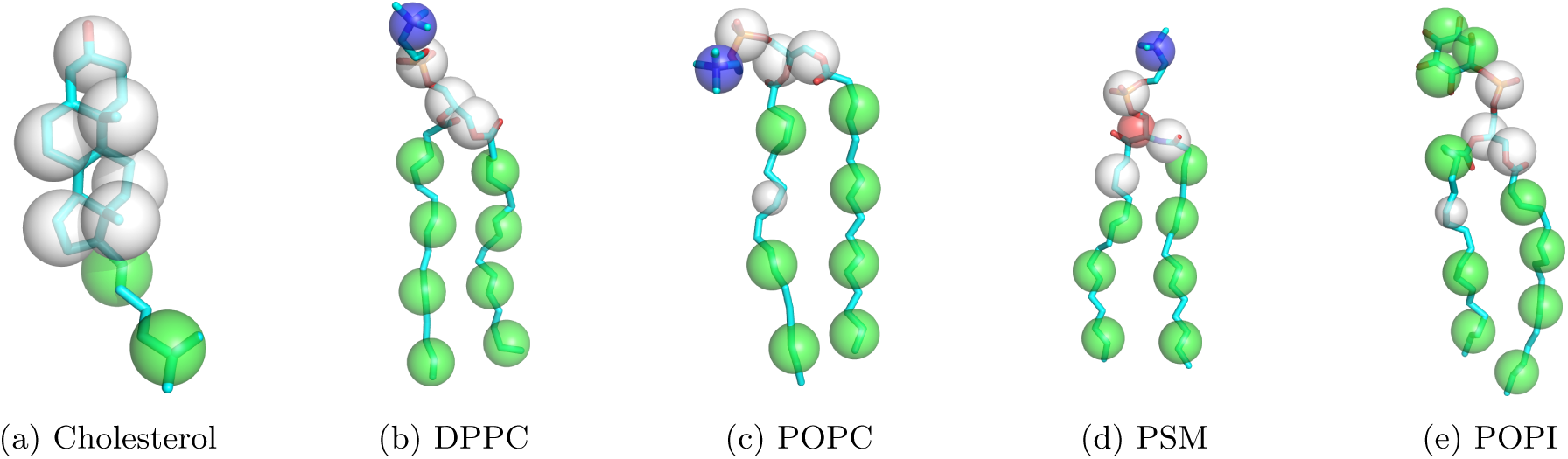
Forward-mapped coarse-grained representations. overlaid on the corresponding all-atom structure, generated with PyCGTOOL. Hydrogens are removed for clarity.

Cholesterol required an additional modification of the molecular graph. Several beads in the Martini 3 cholesterol topology are virtual sites whose positions are defined by virtual-site construction rules and that therefore lack bonded connections to neighboring beads. Because the GNN propagates information exclusively along graph edges, these disconnected virtual-site beads receive no local structural information during message passing, which resulted in distortions of the predicted heavy-atom geometry in the fused-ring region. For the cholesterol input graph, the connectivity was therefore augmented by explicitly connecting each virtual-site bead to its nearest neighboring beads, ensuring that all beads participate in message passing. These additional edges were used only for GNN message passing and did not modify the underlying Martini topology.

Virtual sites are not restricted to cholesterol, and whether they require this treatment depends on whether heavy atoms are assigned to them in the mapping. In the phosphatidylinositol lipids, the virtual bead lies at the center of the inositol ring, where the surrounding beads already account for all ring atoms. No atoms are assigned to the virtual site and it does not enter the molecular graph. In cholesterol, by contrast, the virtual sites lie within the fused-ring system at positions where heavy atoms must be assigned for the mapping to be complete, and these beads therefore necessarily appear in the graph. A virtual-site bead that carries mapped heavy atoms but has no bonded partners consequently requires additional edges, whereas one without mapped atoms is simply omitted. Atom-carrying virtual sites in lipid classes other than sterols were not encountered in the systems studied here and remain to be tested.

### 2.2 Graph representation and feature extraction

Each CG lipid molecule is represented as a graph in which Martini beads form the nodes and every bonded bead pair contributes a pair of directed edges. The bead connectivity is read from per-species connectivity files, which are derived from the Martini 3 topology of each lipid. Node features are derived once per lipid species from the mapping and connectivity files and from the corresponding atomistic CHARMM36 .itp topology files (GROMACS format [18, 19]), and subsequently combined with the bead coordinates for each simulation frame.

Node features comprise: (i) the Martini bead class (categorical, *{C, N, P, Q, W, X, D,* UNK*}*) and bead size (categorical, *{R, S, T,* UNK*}*), both parsed from the bead-type string and passed through lookup-table embeddings; (ii) the Martini bead polarity level, included as a scalar feature; (iii) the bead’s degree in the bonded bead graph (neighbor count); and (iv) the number of heavy atoms mapped to the bead. The latter provides information on the number of atomistic coordinates that must be reconstructed from a given bead and determines the padding mask applied to the network output layer.

The bond, angle, and torsion index tables required by the physics-informed loss terms (Section 2.5) are extracted once per lipid species from the CHARMM topology and cached as lookup templates, together with the ordered list of heavy-atom names defined by the mapping file. Each lipid residue in each simulation frame constitutes one training sample and is stored as a PyTorch Geometric Data object. Each sample contains the node-feature matrix, bead coordinates relative to the lipid center of geometry, target heavy-atom coordinates expressed as displacements from their parent beads, the Boolean output mask, and the auxiliary bond, angle, and torsion index tensors.

The mapping, connectivity and CHARMM36 topology files are per-lipid species definitions rather than per-system inputs, and are required both for training and for backmapping. They define which heavy atoms belong to each bead, the bead graph, and the atom ordering, hydrogen-placement rules and stereochemical targets used to assemble the final atomistic structure. Training additionally requires paired atomistic and forward-mapped coarse-grained trajectories, from which the target displacement vectors are obtained. Backmapping a configuration thus requires only its coarse-grained coordinate file together with the per-lipid species map definitions, which are distributed with MemBack.

### 2.3 Equivariant graph neural network architecture

The CG molecular graphs are processed by an SE(3)-equivariant graph neural network based on the PaiNN architecture [20]. For each bead, the network predicts up to six displacement vectors specifying the positions of the associated heavy atoms relative to the bead center. Geometric information enters the network exclusively through relative position vectors between bonded beads and their distances. Consequently, the predicted displacement vectors are invariant to translations of the input structure and transform equivariantly under rotations; the reconstructed atomistic coordinates therefore transform consistently with the input CG configuration.

Each node carries two parallel representations: an invariant scalar state *s ∈* R*^h^* and an equivariant vector state *v ∈* R*^h×^*^3^, where *h* denotes the hidden dimension. The categorical bead-class and bead-size features are represented by 4and 3-dimensional embeddings, respectively, and concatenated with the scalar polarity, bead-degree, and mapped-heavy-atom-count features (i.e., the local density). The resulting feature vector is linearly projected onto the initial scalar state *s*, while the vector state *v* is initialized to zero. For each bonded edge, the relative displacement vector between the connected beads is computed from their coordinates. Its norm is expanded in a 20-component Gaussian radial basis modulated by a cosine cutoff envelope with a cutoff radius of 12 Å, while the corresponding unit vector provides directional information for equivariant message construction.

Each of the *n* message-passing layers consists of a message block followed by an update block. In the message block, edge-dependent scalar gates are computed from the source-node scalar state and radial-basis representation using two-layer multilayer perceptrons (MLPs). These gates control updates to the scalar and vector states of the destination node. The vector message combines contributions from the source-node vector features, the unit vector along the connecting edge, and their cross product. The cross-product term introduces a parity-sensitive axial-vector contribution, allowing the network to distinguish mirror-related local geometries and thereby encode information required to reconstruct stereochemical features such as chiral centers and cis/trans double bonds.

The subsequent update block applies two equivariant linear transformations to the vector state and constructs rotation-invariant scalar descriptors from vector norms and inner products. Together with the current scalar state, these descriptors are used to compute residual updates to both *s* and *v*. Layer normalization is applied to the scalar channel. Dropout is applied between message-passing layers.

Finally, a gated equivariant readout maps the final vector representation to six vector-valued output channels per bead. Each output vector is modulated by an invariant scalar gate derived from the final scalar state, yielding a 6 *×* 3 array of predicted heavy-atom displacement vectors for each bead. For beads associated with fewer than six heavy atoms, unused output channels are excluded using the padding mask derived from the mapped heavy-atom count. The node representation and overall network architecture are illustrated in Figure 3.

**Figure 3:**
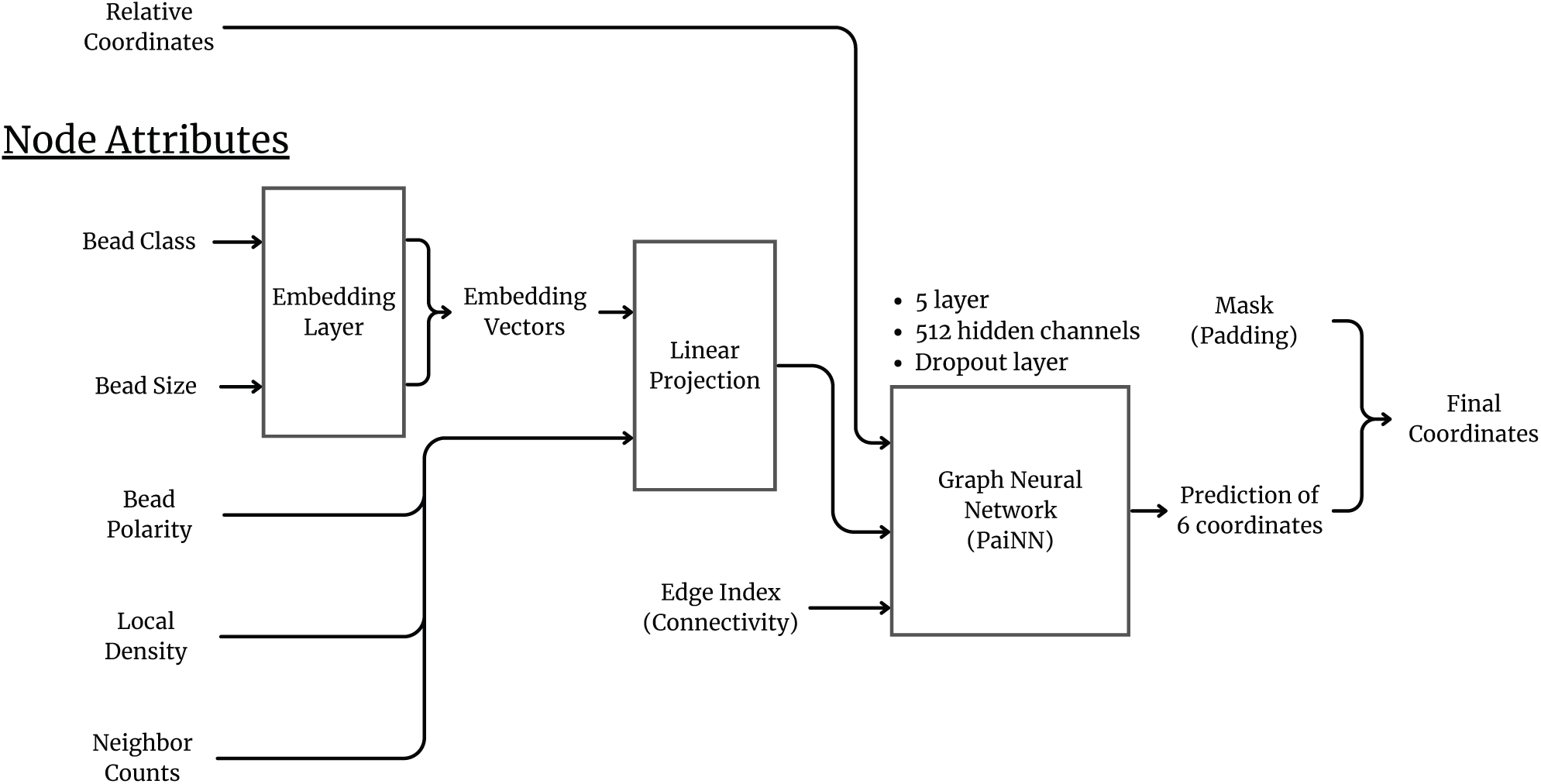
Node features. used by the equivariant graph neural network.

At inference time, the predicted per-bead displacement vectors are added to the corresponding CG bead position and offset by the lipid’s center of geometry to obtain absolute heavy-atom coordinates, and the resulting atoms are used, together with the mapping file, to construct an all-heavy-atom system.

### 2.4 Training simulation database and data pipeline

The training database comprises 37 atomistic lipid-bilayer trajectories together with their corresponding forward-mapped Martini 3 representations. The dataset includes e.g. a PSM:POPC:cholesterol bilayer, pure DPPC and DMPC bilayers, a multicomponent red-blood-cell (RBC) membrane model, and a broader library of bilayers containing phosphatidic acid (PA), phosphatidylethanolamine (PE), phosphatidylserine (PS), phosphatidylglycerol (PG), phosphatidylinositol (PI), phosphatidylcholine (PC), sphingomyelin (SM), and cardiolipin (CL), either individually or in multicomponent mixtures and with or without cholesterol. Simulations were performed at temperatures between 300 K and 350 K at an ion concentration of 0.15 M NaCl. The complete set of training systems is summarized in the Supporting Information, Table S1. In total, the database contains 2,691,038 lipid graphs covering 50 distinct lipid species. These were drawn from a pool of roughly 58 µs of atomistic trajectory data. Frames were not sampled at a uniform interval across systems: depending on trajectory length and the number of lipids per frame, either every frame or every *n*-th frame was used, with sampling intervals ranging from 1 to 40 ns. Systems contributing fewer lipids per frame were sampled more densely, so that no single large multicomponent system dominates the dataset.

For each atomistic/CG trajectory pair, bead and atom coordinates are loaded once per trajectory, after which construction of the per-residue training graphs is parallelized across CPU worker processes. For each lipid and simulation frame, the corresponding heavy-atom displacement vectors relative to their parent beads are generated as training targets. This parallelized pipeline reduced the wall-clock time required to process 100 ns of trajectory data for a multicomponent system from approximately 1.5 h to 13 s. Frames were sampled such that consecutive frames entering the database are separated by at least 1 ns of simulation time. The resulting graphs are split frame-wise into training, testing, and validation subsets in a ratio of 70:20:10 using a fixed random seed. All lipid graphs originating from a given frame are assigned to the same subset, so that lipids of one configuration are never distributed across subsets.

Because the complete training dataset exceeds available GPU memory, samples are pre-batched into groups of 1024 graphs (2,630 batches in total) and stored in an LMDB key-value database. Training, test, and validation subsets are maintained as separate LMDB environments, allowing the database to be extended incrementally as additional simulation data become available. During training, a PyTorch DataLoader streams the pre-batched samples using multiple worker processes, pinned memory, persistent workers, and a prefetch factor of 8. A CUDA-stream-based prefetcher additionally overlaps host-to-device transfer of the subsequent batch with GPU computation on the current batch, thereby minimizing data-loading overhead during training.

### 2.5 Model training

The network was trained by minimizing a weighted, physics-informed loss evaluated on the reconstructed heavy-atom coordinates. Given the predicted and reference heavy-atom positions, the objective comprises four terms: (i) a coordinate-distance term, defined as the superposition-free root-mean-square deviation between predicted and reference atomic positions; (ii) a bond-length term, defined as the mean squared error (MSE) between predicted and reference bond lengths for all heavy-atom bonds specified by the CHARMM topology; (iii) an angle term, defined as the MSE between predicted and reference bond angles, calculated in radians for all bonded atom triplets; and (iv) a torsion term that penalizes deviations in a selected set of stereochemically relevant dihedral angles (R/S chirality, cis-trans conformations at acyl chain and gauche conformations at ring structures). For the torsion term, angular differences are evaluated using an atan2-based formulation and wrapped to the interval [*−π, π*] to preserve periodicity. Dihedrals for which the angle is numerically ill-defined because of near-collinear bond vectors are excluded from both the loss and gradient calculation. The four included terms are combined as a weighted sum, with a larger relative weight placed on the bond, angle, and torsion terms than on the coordinate-distance term, to favor chemically consistent local geometries over a purely coordinate-based reconstruction (weights: (*w*_coord_*, w*_bond_*, w*_angle_*, w*_torsion_) = (1, 5, 5, 5)).

Optimization was performed using Adaptive Moment Estimation (Adam) [21] with an initial learning rate of 4 *×* 10*^−^*^4^ and gradient-norm clipping at 10. A ReduceLROnPlateau scheduler reduced the learning rate by a factor of two when the validation loss failed to improve by at least 1% over four consecutive epochs, with a minimum learning rate of 1 *×* 10*^−^*^7^. Training was terminated early if the validation loss failed to improve by at least 0.5% over 15 consecutive epochs, and the model checkpoint with the lowest validation loss was retained for subsequent analyses. The production model comprised five message-passing/update layers with a hidden dimension of 512 and a dropout probability of 0.04. These configurations were selected by hyperparameter search based exclusively on performance on the validation subset.

### 2.6 Hydrogen reconstruction

Because the GNN predicts heavy-atom coordinates only, hydrogen atoms are added in a separate post-processing step. Several hydrogen-placement strategies were evaluated. The pdb2gmx routine implemented in GROMACS produced the most reliable hydrogen geometries but is primarily designed to generate complete molecular topologies and did not preserve the lipid-specific residue definitions required by the subsequent workflow. We therefore reimplemented the hydrogen-placement procedure used by pdb2gmx in Python. Hydrogen coordinates are generated using the same lipid-specific hydrogen database (.hdb) entries employed by GROMACS, which define hydrogen placement from the local geometry of the corresponding heavy atom and its bonded neighbors.

### 2.7 Water and ion conversion

CG water and ion beads are converted to their atomistic representations before energy minimization. Each standard Martini 3 water bead is replaced by four explicit TIP3P water molecules. Their oxygen atoms are positioned at tetrahedrally distributed offsets from the CG bead center, with the offset distance chosen such that all oxygen atoms remain within the volume represented by the original CG bead. Each water molecule is subsequently assigned an independent random orientation, sampled uniformly in three-dimensional rotational space while keeping its oxygen position fixed. The resulting coordinates are wrapped into the primary simulation cell according to the periodic boundary conditions. Ion beads are converted one-to-one to the corresponding atomistic ion residues according to a predefined lookup table, with the atomistic ion initially placed at the position of its parent CG bead. This one-to-one correspondence follows from the Martini 3 ion model.

### 2.8 Clash detection and resolution

Because each Martini bead represents several heavy atoms, a non-overlapping CG configuration does not necessarily remain free of steric clashes after reconstruction at atomistic resolution. This problem occurs particularly in densely packed regions of the bilayer, where beads from neighboring lipids or opposing lipid tails may be sufficiently close that their reconstructed heavy atoms overlap. The effect can be amplified by variations in mapping multiplicity along a lipid chain: while most beads represent four heavy atoms, some mappings assign a larger number, producing correspondingly extended atomistic segments. Consequently, CG bead separations that are unproblematic at coarse-grained resolution can yield sterically overlapping atomistic configurations after backmapping. Such clashes were observed particularly in complex multicomponent systems and motivated the additional clash-resolution procedure described below.

Additional steric overlaps can arise during subsequent hydrogen reconstruction. Clashing atom pairs are identified using a periodic neighbor search with a predefined distance threshold (0.3 Å). Clash resolution is performed iteratively, starting with the atom pairs exhibiting the smallest interatomic distances. For each selected pair, the two atoms are displaced symmetrically in opposite directions along their periodic minimum-image separation vector until a threshold distance (0.3 Å) is reached. To reduce interference between simultaneous corrections, no more than one clash involving a given lipid molecule is resolved within a single iteration. The search and correction steps are repeated until no atom pairs remain below the clash threshold or a predefined maximum number of iterations is reached.

### 2.9 Chirality correction

Although the training objective includes a stereochemistry-sensitive torsional term (Section 2.5), a small fraction of stereocenters are reconstructed with inverted configuration. These inversions are corrected deterministically during post-processing. Target stereochemistry is obtained from the improper-dihedral restraints defined in the GROMACS CHARMM .itp topology. For the relevant stereocenters, equilibrium improper-dihedral values of *±*120*^◦^* encode the required configuration (*R/S*, depending on atom ordering).

For each stereocenter, the handedness of the reconstructed geometry is determined from the sign of the scalar triple product formed by the vectors from the central atom to its three heavy-atom substituents. If this sign is inconsistent with the configuration specified by the topology, the stereocenter is inverted geometrically. The central atom and its attached hydrogen are reflected across the plane defined by the three heavy-atom substituents, while the substituent atoms themselves remain fixed. Because reflection preserves the distances from the central atom to the three atoms defining the plane, this operation reverses the local handedness while leaving the positions of the surrounding heavy-atom framework unchanged.

Formally, for a stereocenter *c* with heavy substituents *a*, *b*, *d* and hydrogen *H*, the handedness is given by the sign of the scalar triple product

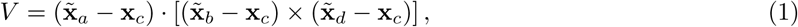

where **x̃** denotes the minimum-image position relative to **x***_c_*. The target sign *s^∗^* is obtained from the sign of the corresponding *±*120*^◦^* improper-dihedral restraint in the topology. If sign(*V*) = *s^∗^*, the center is left unchanged. Otherwise, let

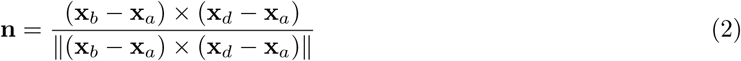

be the unit normal of the plane through *a*, *b* and *d*. The center and its hydrogen are then reflected across this plane,

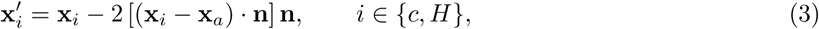

while all remaining atoms are left unchanged (Figure 4).

**Figure 4:**
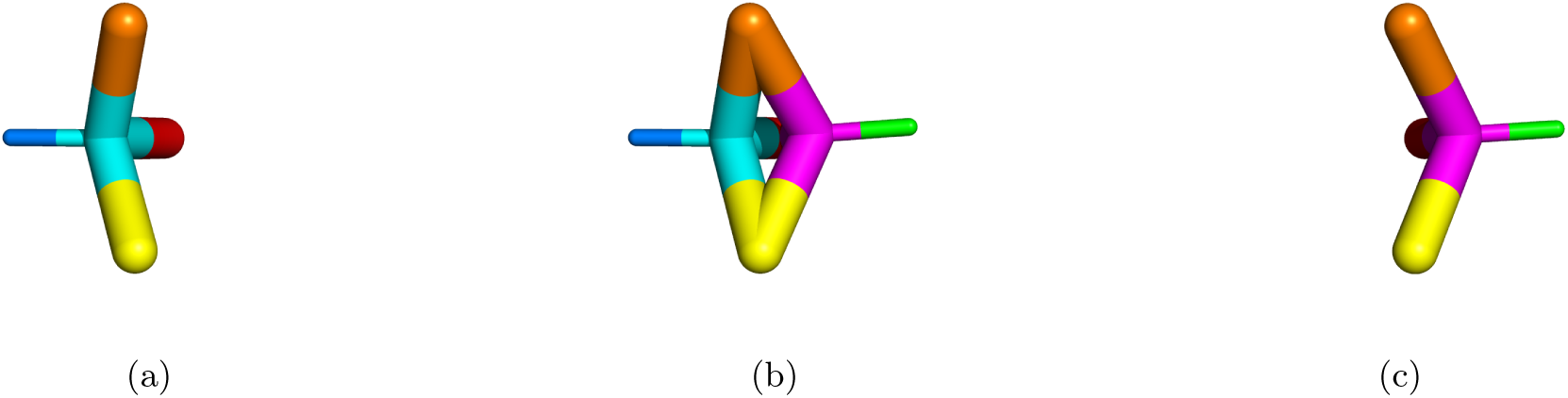
Geometric correction of an inverted stereocenter. Shown is the glycerol *C*2 stereocenter of a DPPC molecule whose configuration was reconstructed with inverted handedness. (a) Reconstructed geometry, with the central atom *C*2 (cyan), its three heavy substituents *C*1 (orange), *C*3 (yellow) and *O*21 (red), which define the reflection plane, and the attached hydrogen *HS* (blue). (b) Superposition of the reconstructed (cyan) and corrected (magenta) geometries. The three heavy substituents lie in the reflection plane and are common to both. (c) Corrected geometry after reflection of *C*2 (magenta) and *HS* (green) across that plane. Bond lengths and bond angles at the stereocenter are preserved exactly, and no substituent atom is displaced.

Because *a*, *b* and *d* lie in the reflection plane, they are fixed points of the transformation. The reflection is an isometry with determinant *−*1, so all distances **x***_c_ −* **x***_k_* and angles ∠(*k, c, l*) for *k, l ∈ {a, b, d, H}* are preserved exactly, whereas *V ^′^* = *−V* . The operation therefore inverts the local handedness without altering any bond length or bond angle at the stereocenter, and without displacing the substituents or the molecular fragments attached to them. Since one of the four substituents must be a hydrogen for the reflection to be defined in this way, stereocenters bearing four heavy substituents are not processed; no such center occurs among the restrained stereocenters of the lipid topologies used here.

### 2.10 System assembly and molecular dynamics protocol

For each backmapped system, a simulation-ready input set is generated automatically. The CHARMM .itp topology files corresponding to the lipid species present in the reconstructed system are collected in a toppar directory together with the required force-field files, and a system topology (topol.top) is generated according to the residue composition of the reconstructed structure. User-provided topology files can override entries in the internal topology database, allowing the workflow to accommodate lipid species not included by default.

The reconstructed system is first subjected to steepest-descent energy minimization to a maximum-force tolerance of 1, 000 kJ mol*^−^*^1^ nm*^−^*^1^. Non-bonded interactions are evaluated using Verlet neighbor lists, force-switched van der Waals interactions with switching between 1.0 and 1.2 nm, and particle-mesh Ewald (PME) electrostatics with a real-space cutoff of 1.2 nm. During minimization, positional restraints with a force constant of 1, 000 kJ mol*^−^*^1^ nm*^−^*^2^ are applied to headgroup atoms of the lipids, while dihedral restraints of 1, 000 kJ mol*^−^*^1^ rad*^−^*^2^ are imposed on stereochemically relevant centers to preserve the reconstructed geometry and the corrected stereochemistry.

Equilibration follows a six-stage protocol adapted from the standard CHARMM-GUI [22] membrane equilibration scheme and spans a total of 1.875 ns (Supporting Information, Table S2). Positional and dihedral restraints are progressively reduced across the equilibration stages, allowing the reconstructed atomistic structure to relax gradually while avoiding large atomic displacements during the initial dynamics. The initial stages use a 1 fs integration time step, followed by 2 fs in the later stages. Temperature is maintained at 303.15 K by velocity-rescaling coupling, while semi-isotropic C-rescale pressure coupling [23] is introduced during the NPT stages. All restraints are removed in the final equilibration stage. Subsequent production simulations are performed without positional or stereochemical restraints.

### 2.11 Software development and availability

The feature-extraction, model-training, and post-processing components of *MemBack* are maintained under version control on GitHub, with a continuous integration/continuous deployment (CI/CD) pipeline that runs an accompanying unit test suite and builds project documentation on each change, and the package is distributed for installation via PyPI. Trajectory handling and analysis use MDAnalysis [24, 25]. Proposed software is publicly accessible at *MemBack* GitHub repository.

## 3 Results and discussion

### 3.1 Model training and convergence

The production model was trained using the architecture and optimization protocol described in Section 2.5. Each epoch comprised 1840 pre-batched training batches and required approximately 268–278 s on a single NVIDIA RTX 5090 GPU, corresponding to a total training time of 278.3 min. The model consists of *≈*16 million trainable parameters. The pre-batched LMDB data pipeline rendered data-loading overhead negligible compared with the GPU computation time.

Training converged rapidly during the initial epochs (Supporting Information, Figure S1). The validation loss decreased from 1.359 after the first epoch to 0.870 at epoch 10, while the mean per-lipid coordinate RMSD decreased from 0.92 Å to 0.73 Å. This initial phase was followed by a slower improvement, with the validation loss reaching approximately 0.80 and the coordinate RMSD approximately 0.68 Å by epoch 35. Subsequent changes were small. The learning-rate scheduler was first activated at epoch 19 and progressively reduced the learning rate from 4 *×* 10*^−^*^4^ to its minimum value of 2 *×* 10*^−^*^6^. Training terminated by early stopping at epoch 61, and the checkpoint from epoch 46 was retained according to the validation-loss criterion.

At the retained checkpoint, the training and validation losses were 0.802 and 0.795, respectively, with corresponding coordinate RMSDs of 0.68 Å and 0.67 Å. The bond, angle, and torsion components likewise showed closely matched values for the training and validation sets (0.07 Å/0.07 Å, 0.08/0.08, and 0.11/0.11, respectively; training/validation). The close agreement between the two datasets throughout optimization provides no indication of appreciable overfitting under the present data split. Instead, both curves approach a common plateau, indicating that further optimization with the selected architecture and loss formulation yields only limited improvement. Whether this residual error reflects model capacity, optimization constraints, or the intrinsic structural degeneracy of the coarse-grained representation cannot be distinguished from the learning curves alone.

### 3.2 Reconstruction accuracy

We first quantified reconstruction accuracy for configurations for which the corresponding atomistic structures were available as a reference. Each benchmark frame was taken from a trajectory segment excluded from model training, whereas earlier frames from the same trajectory contributed to the training database. The reference atomistic configuration was forward-mapped to its Martini 3 representation, reconstructed with MemBack and minimized (see Section 2.10), and compared directly with the original atomistic coordinates. This benchmark therefore assesses reconstruction of previously unseen configurations while retaining lipid chemistries and membrane systems represented during training.

Throughout this section, RMSD refers to the superposition-free per-lipid heavy-atom RMSD. Predicted and reference coordinates are compared directly using the periodic minimum-image convention, without rigidbody alignment. This definition differs from the optimally superposed RMSD commonly used in structural biology. The distinction is important for backmapping: the CG coordinates constrain the spatial position and orientation of each lipid, and errors in either should therefore contribute to the reconstruction metric rather than being removed by post hoc alignment.

Table 1 summarizes the reconstruction accuracy across 15 benchmark systems. For forward-mapped inputs, MemBack achieved a mean per-lipid RMSD of 0.65 Å, with system-specific values ranging from

**Table 1:** Reconstruction accuracy for held-out trajectory frames and held-out membranes. Each row corresponds to a single atomistic frame taken from a trajectory segment excluded from model training, although earlier frames from the same trajectory were included in the training database, or to frames of a membrane system not included in model training (marked by *‡*). *^∗^*, lipid species absent from the training database. RMSD is the superposition-free per-lipid heavy-atom RMSD and is reported as mean *±* standard deviation over all lipids in the frame. “FM” denotes direct forward-mapped CG input. The Backward column reports the initial geometric projection of that tool, i.e. the structure obtained before its subsequent energy-minimization and restrained-MD relaxation, and serves as a baseline for the accuracy attainable without a learned or optimized component. The MemBack “FM, projection” column reports the same MemBack reconstructions before energy minimization, for direct comparison with the Backward projection. Additionally, bond, angle, and dihedral errors are reported for the minimized MemBack structures obtained from direct forward-mapped inputs. Reconstruction times denote wall-clock time per frame. Times in the parenthesis are times for minimization. Mean values are pooled over all lipids of all benchmark systems.

| System | MemBack RMSD (Å) |  | CG2AT2 RMSD (Å) | Backward RMSD (Å) | MemBack geometry (FM) |  |  | MemBack | CG2AT2 |
| --- | --- | --- | --- | --- | --- | --- | --- | --- | --- |
|  | FM | FM, projection | FM | FM, projection | Bond (Å) | Angle (°) | Dihed. (°) | Time (seconds) |  |
| PSM:POPC:Chol | 0.64 ± 0.23 | 0.62 ± 0.21 | 1.09 ± 0.20 | 1.42 ± 0.26 | 0.03 | 3.8 | 6.1 | 7.87 (4) | 30 (7) |
| DPPC | 0.89 ± 0.11 | 0.83 ± 0.11 | 1.23 ± 0.11 | 1.44 ± 0.05 | 0.04 | 4.1 | 4.8 | 1.33 (1) | 5 (1) |
| DMPA:DMPE | 0.54 ± 0.18 | 0.53 ± 0.17 | 0.95 ± 0.14 | 1.57 ± 0.05 | 0.03 | 3.8 | 9.7 | 1.53 (1) | 8 (2) |
| DOPE:DOPS | 0.71 ± 0.12 | 0.69 ± 0.12 | 1.11 ± 0.11 | 1.83 ± 0.07 | 0.03 | 4.5 | 6.2 | 1.47 (1) | 10 (3) |
| DOPG:DOPE | 0.70 ± 0.14 | 0.68 ± 0.13 | 1.08 ± 0.11 | 1.83 ± 0.08 | 0.03 | 4.3 | 6.3 | 1.58 (1) | 8 (2) |
| PLPG:PLPE | 0.77 ± 0.12 | 0.73 ± 0.11 | 1.03 ± 0.12 | 1.62 ± 0.09 | 0.03 | 4.3 | 6.3 | 1.50 (1) | 9 (3) |
| POPA | 0.76 ± 0.08 | 0.73 ± 0.08 | 1.07 ± 0.11 | 1.66 ± 0.06 | 0.03 | 4.2 | 5.8 | 1.49 (1) | 8 (1) |
| POPI:POPE | 0.59 ± 0.13 | 0.58 ± 0.12 | 0.97 ± 0.11 | 1.64 ± 0.06 | 0.03 | 4.2 | 6.1 | 1.56 (1) | 10 (3) |
| POPI33:POPE:POPS:Chol | 0.56 ± 0.20 | 0.54 ± 0.19 | 0.99 ± 0.17 | 1.46 ± 0.27 | 0.03 | 4.6 | 6.2 | 1.66 (1) | 12 (2) |
| SAPI:POPE:Chol | 0.52 ± 0.17 | 0.50 ± 0.16 | 0.93 ± 0.14 | 1.50 ± 0.29 | 0.03 | 4.0 | 6.6 | 1.63 (1) | 11 (3) |
| SDPA:SDPE | 0.73 ± 0.13 | 0.71 ± 0.12 | 1.12 ± 0.11 | 1.45 ± 0.07 | 0.03 | 5.3 | 7.0 | 2.50 (2) | 10 (2) |
| SOPA | 0.67 ± 0.14 | 0.65 ± 0.14 | 1.10 ± 0.12 | 1.69 ± 0.06 | 0.04 | 4.4 | 6.2 | 2.61 (2) | 7 (1) |
| TOCL2:POPE:POPC | 0.70 ± 0.14 | 0.68 ± 0.13 | 1.07 ± 0.11 | 1.61 ± 0.10 | 0.03 | 4.1 | 6.1 | 2.52 (2) | 10 (3) |
| DPPC:DLIPC:Chol<sup>†</sup> | 0.71 ± 0.31 | 0.71 ± 0.32 | 1.07 ± 0.21 | 1.42 ± 0.34 | 0.03 | 5.0 | 6.8 | 2.75 (2) | 10 (3) |
| SCD Red Blood Cell<sup>‡</sup> [26] | 0.60 ± 0.28 | 0.58 ± 0.26 | 0.97 ± 0.18 | 1.36 ± 0.32 | 0.03 | 4.0 | 7.8 | 6.83 (3) | 29 (6) |
| DLIPC* | 0.96 ± 0.10 | 1.03 ± 0.08 | — | — | 0.04 | 6.7 | 7.0 | — | — |
| PDOPE* | 0.74 ± 0.11 | 0.71 ± 0.10 | — | — | 0.03 | 5.3 | 7.7 | — | — |
| PLA20* | 0.89 ± 0.08 | 0.84 ± 0.07 | — | — | 0.04 | 5.8 | 7.8 | — | — |
| Mean | 0.65 Å | 0.62 Å | 1.04 Å | 1.49 Å | 0.03 Å | 4.3° | 6.5° | 2.59 s | 11.80 s |

0.52 Å for SAPI:POPE:Chol to 0.89 Å for DPPC. Applied to the same forward-mapped configurations, CG2AT2 yielded a mean RMSD of 1.04 Å using its fragment-fitting and minimization workflow. MemBack yielded a lower mean RMSD for all 15 systems examined, and the pooled per-lipid distributions show a pronounced shift toward lower reconstruction errors (Figure 5). To assess the reconstruction accuracy prior to minimization, we additionally compared MemBack (mean RMSD: 0.62 Å before minimization) with the purely geometric projection used by Backward [6] (RMSD: 1.49 Å before minimization; Table 1). Whereas the geometric projection recovers only the approximate atom positions, MemBack reconstructs their atomistic arrangement, reducing the mean RMSD compared to geometric projection by more than a factor of two.

**Figure 5:**
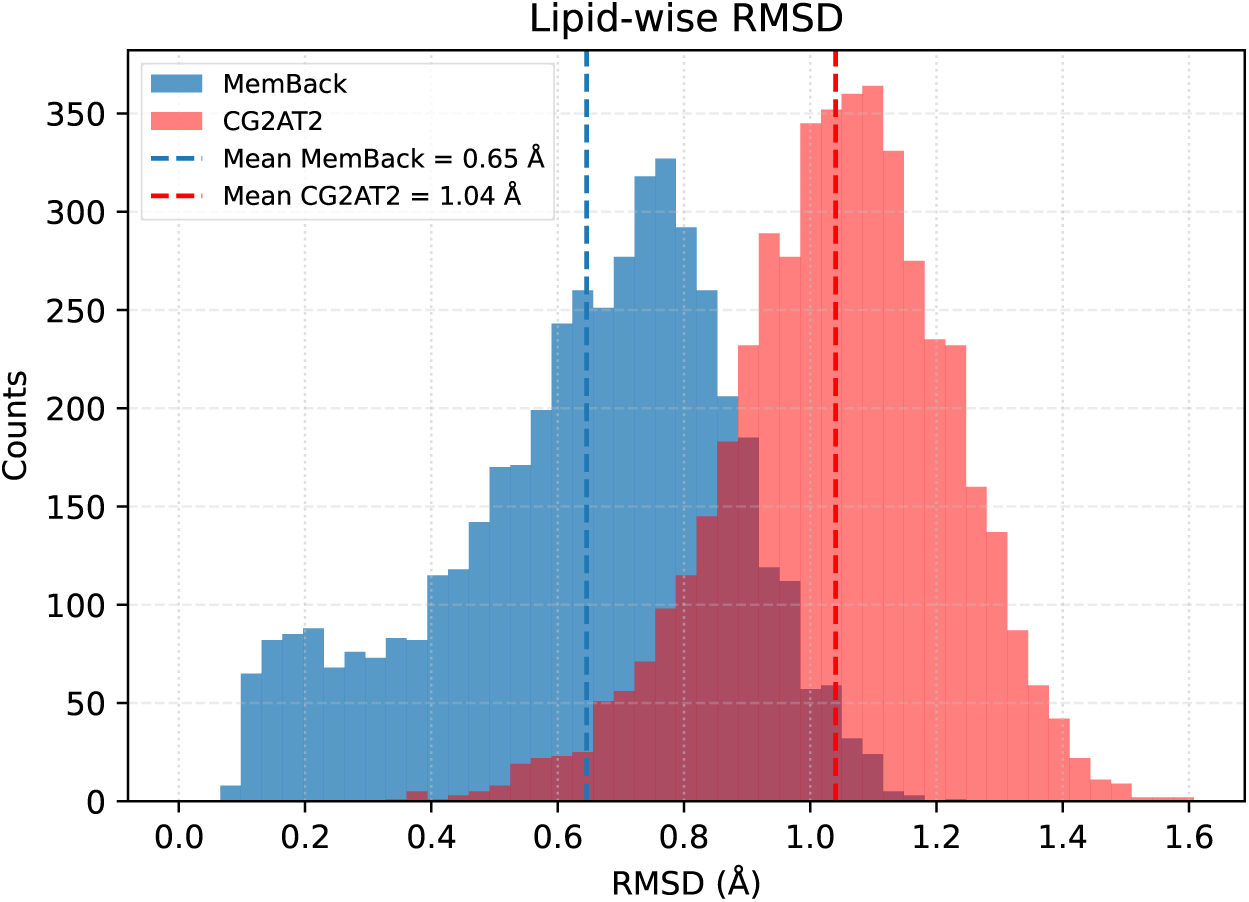
Pooled distributions of the superposition-free per-lipid heavy-atom RMSD for MemBack and CG2AT2 across all benchmark systems. The mean RMSD is 0.65 Å for MemBack and 1.04 Å for CG2AT2.

Table 1 summarizes the reconstruction accuracy across 15 benchmark systems. For forward-mapped inputs, MemBack achieved a mean per-lipid RMSD of 0.65 Å, with system-specific values ranging from 0.52 Å for SAPI:POPE:Chol to 0.89 Å for DPPC. Applied to the same forward-mapped configurations, CG2AT2 yielded a mean RMSD of 1.04 Å using its fragment-fitting and minimization workflow. MemBack yielded a lower mean RMSD for all 15 systems examined, and the pooled per-lipid distributions show a pronounced shift toward lower reconstruction errors (Figure 5). To place these values on an absolute scale, we additionally compared the mean RMSD before minimization (0.62 Å) with the purely geometric projection performed by Backward [6], which yielded a mean per-lipid RMSD of 1.49 Å (before minimization and equilibration, Table 1). Projecting atoms onto their parent bead positions recovers only the approximate location of each atom, and the learned reconstruction reduces the RMSD by more than a factor of two.

To probe reconstruction beyond membrane systems represented during training, the benchmark set included two membrane models that were entirely excluded from the training database (marked by *‡* in Table 1): a DPPC:DLIPC:Chol bilayer and a red-blood-cell (RBC) membrane model [26]. We refer to this held-out membrane as the symmetric cholesterol distribution (SCD) model, and to the RBC membrane contained in the training database [27] (Table S1) as the asymmetric cholesterol distribution (ACD) model. The two RBC plasma membrane models were derived from different lipidomics and biophysical analyses and differ substantially in their transbilayer organization (see [28] for in-depth comparison of SCD and ACD models). The SCD model has comparable phospholipid numbers and a symmetric cholesterol distribution in its two leaflets, whereas the ACD model combines an approximately 30% phospholipid excess in the cytoplasmic leaflet with a strongly asymmetric cholesterol distribution. A direct comparison of the two models, including their lipid compositions, leaflet organization, cholesterol distribution, and which species entered the training database, is given in the Supporting Information (Table S5). The two excluded membrane systems also contain lipid species that were not represented during training, providing an additional test of transfer beyond the specific molecular compositions encountered by the model.

Despite these differences, MemBack achieved mean per-lipid RMSDs of 0.71 Å for the DPPC:DLIPC:Chol bilayer and 0.60 Å for the RBC membrane. These values are comparable to those obtained for membrane systems represented during training, indicating that reconstruction accuracy is retained for previously unseen membrane compositions and organizations. Transfer to the individual lipid species absent from the training database is examined in more detail below.

MemBack also substantially reduced the computational cost of reconstruction. Across the benchmark systems, reconstruction required 2.59 s per frame on average, compared with 11.80 s for CG2AT2, corresponding to an approximately 4-fold reduction in wall-clock time under the benchmark conditions used here.

### 3.3 Generalization to unseen lipid species

The benchmarks above establish that MemBack reconstructs previously unseen configurations with high accuracy and further indicate transfer to membrane compositions excluded from training. We next examined whether this transfer extends to individual lipid species that were not encountered during model training. Importantly, lipid residue identity is not provided to the network as an input feature. Atomistic reconstruction is instead conditioned on Martini bead types, bonded connectivity, and local molecular geometry. The representation should therefore permit transfer to previously unseen lipid molecules when their constituent bead types and local bonded motifs are represented within the training data.

DLIPC provides one such test. This lipid was absent from the training database but is present in the DPPC:DLIPC:Chol benchmark system. For a forward-mapped reference configuration, MemBack reconstructed DLIPC with a mean per-lipid heavy-atom RMSD of 0.96 Å, without lipid-specific retraining or modification of the reconstruction procedure. The corresponding native Martini 3 DPPC:DLIPC:Chol membrane could likewise be backmapped, minimized, equilibrated, and propagated in atomistic MD without species-specific intervention.

Transfer was further quantified for PLA20 and PDOPE in the RBC benchmark membrane. Both lipid species were excluded from the training database but were present in the forward-mapped reference configuration. MemBack reconstructed PLA20 and PDOPE with mean per-lipid heavy-atom RMSDs of 0.89 Å and 0.74 Å, respectively. These reconstruction errors are comparable to those obtained for lipid species represented during training in the same membrane system. Moreover, the corresponding native Martini 3 membrane could be backmapped, minimized, equilibrated, and propagated in an unrestrained atomistic simulation without lipid-specific intervention.

These results show that successful reconstruction does not require the complete lipid molecule to have been encountered during training. Rather, the model can combine learned local bead-level environments to reconstruct previously unseen lipid species assembled from familiar Martini bead types and bonded motifs. This form of transfer is particularly relevant for lipid force fields, in which chemically distinct species frequently share recurring headgroup, glycerol-backbone, and acyl-chain building blocks. The present results do not, however, establish extrapolation to novel Martini bead types or fundamentally different local chemical environments, which remain to be tested.

### 3.4 Geometric and stereochemical fidelity

Low coordinate error alone does not ensure chemically valid atomistic structures. We therefore evaluated local molecular geometry independently of the coordinate-based reconstruction error. Across the benchmark systems, MemBack reproduced heavy-atom bond lengths with a mean RMSE of 0.03 Å and bond angles with a mean RMSE of 4.3*^◦^*. The subset of stereochemically relevant dihedral angles used to constrain the reconstruction was reproduced with a mean RMSE of 6.5*^◦^* (Table 1). Thus, the sub-Å coordinate accuracy of the reconstructed lipids is accompanied by preservation of their local bonded geometry. These quantities are reported only for MemBack. For CG2AT2, the quality of the corresponding local geometric terms depends on lipid-specific mapping optimization, which was not performed here; a direct comparison would therefore not represent the fully optimized performance of that method.

Stereochemical fidelity was assessed separately for tetrahedral stereocenters and configurationally constrained unsaturated lipid double bonds. For each reconstructed lipid, the configuration of every annotated stereocenter and double bond was determined and compared with that of the corresponding atomistic reference structure. Following the deterministic chirality-correction step (Section 2.9) and before energy minimization, 100.0% of stereocenters and 100.0% of double bonds retained the reference configuration for directly forward-mapped inputs. These results show that the combined equivariant reconstruction and stereochemical post-processing pipeline preserves the configurational identity of the reference lipids with near-complete fidelity.

### 3.5 Hydrogen placement

The hydrogen-reconstruction procedure described in Section 2.6 was validated independently of the GNN-based heavy-atom reconstruction using four systems from the simulation database: PSM:POPC:Chol, DMPA: DMPE, pure DPPC, and the multicomponent RBC membrane. For one frame from each system, hydrogen atoms were removed from the reference atomistic structure and subsequently reconstructed from the unchanged heavy-atom coordinates. The rebuilt hydrogen coordinates were then compared directly with the corresponding reference positions. This procedure isolates errors arising from the hydrogen-placement algorithm from those introduced during heavy-atom backmapping (before minimization). Because the reference hydrogens are configurations sampled during molecular dynamics, whereas the reconstruction places them at idealized bond lengths and angles, exact agreement is not expected. The procedure is illustrated for a PSM molecule in the Supporting Information, Figure S3.

Across the four systems, the pooled per-lipid hydrogen atom-position RMSD was 0.34 *±* 0.13 Å (mean *±* standard deviation; Supporting Information, Figure S2). The close agreement with the reference geometry demonstrates that the reimplemented hydrogen-building procedure reproduces the hydrogen atom coordinates with high accuracy and provides suitable starting geometries for subsequent energy minimization.

### 3.6 Membrane observables

Reconstruction accuracy at the level of individual lipids does not by itself establish that a backmapped configuration reproduces the structural characteristics of the bilayer. We therefore examined whether ensemble-averaged membrane observables computed from MemBack reconstructions agree with those of the underlying atomistic reference. The test system was a RBC membrane model composed of 1,168 lipids distributed over 16 species (Chol, DPPC, LSM, NSM, PAPC, PAPS, PDOPE, PLA20, PLPC, POPC, POPE, PSM, SAPE, SAPI2A, SAPS, and SOPC), with cholesterol accounting for 476 molecules, or 40.8 mol% of the membrane, distributed symmetrically between the two leaflets. The simulation box measured 16.03*×*16.03 nm^2^ in lateral direction. The atomistic reference trajectory for this composition was taken from [28].

One hundred configurations were extracted from the atomistic reference trajectory at 10 ns intervals between 2 and 3 µs. Each configuration was forward-mapped to its Martini 3 representation, solvated and neutralized with NaCl using insane.py [29], and subsequently reconstructed with MemBack. Each reconstructed frame was minimized for 100 steps with positional and dihedral restraints applied as described in Section 2.10, and no further equilibration or production dynamics were performed. Each backmapped frame is compared against the atomistic configuration from which it originated. Bilayer thickness, area per lipid (APL), and the acyl-chain order parameter *S_CC_* were computed with liPyphilic [30], and APL and *S_CC_* were pooled over all lipids and all 100 frames. Membrane thickness was computed as the difference between the mean membrane leaflet positions along the membrane normal (*z*), at each frame. Leaflet positions were defined by the phosphorus atoms of the phospholipids and the hydroxyl oxygen of cholesterol, with leaflet membership assigned per frame using LiPyphilic. A single global height per leaflet was used rather than a local intrinsic surface.

Bilayer thickness was tracked frame by frame across the 1 µs interval (Figure 6). The reconstructed membranes reproduce both the magnitude and the frame-to-frame variation of the reference thickness, with all values confined to a window of approximately 41.9 to 42.7 Å. The two traces follow one another across individual frames rather than merely agreeing on average, indicating that the reconstruction preserves the configuration-specific vertical organization of the bilayer.

**Figure 6:**
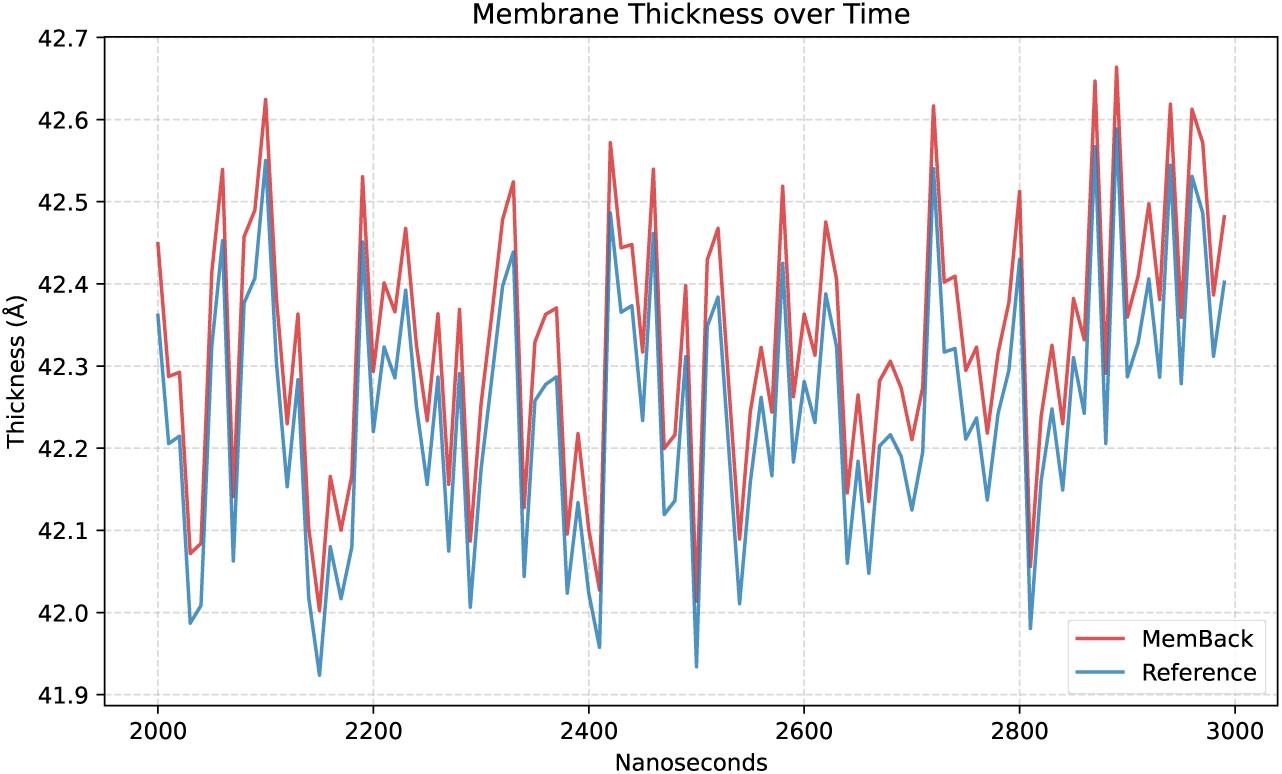
Membrane thickness comparison for 100 backmapped snapshots over 1*µs*. Backmapped results are minimized for 100 steps with position and dihedral restraints.

The pooled APL distributions of the reconstructed and reference membranes are shown in Figure 7a. Both distributions yield an average of 44.44 Å^2^ and overlap over their full range. The shape of the distribution, which reflects how the reconstructed atoms are distributed around those positions, indicates that MemBack does not introduce packing defects or locally over-expanded regions.

**Figure 7:**
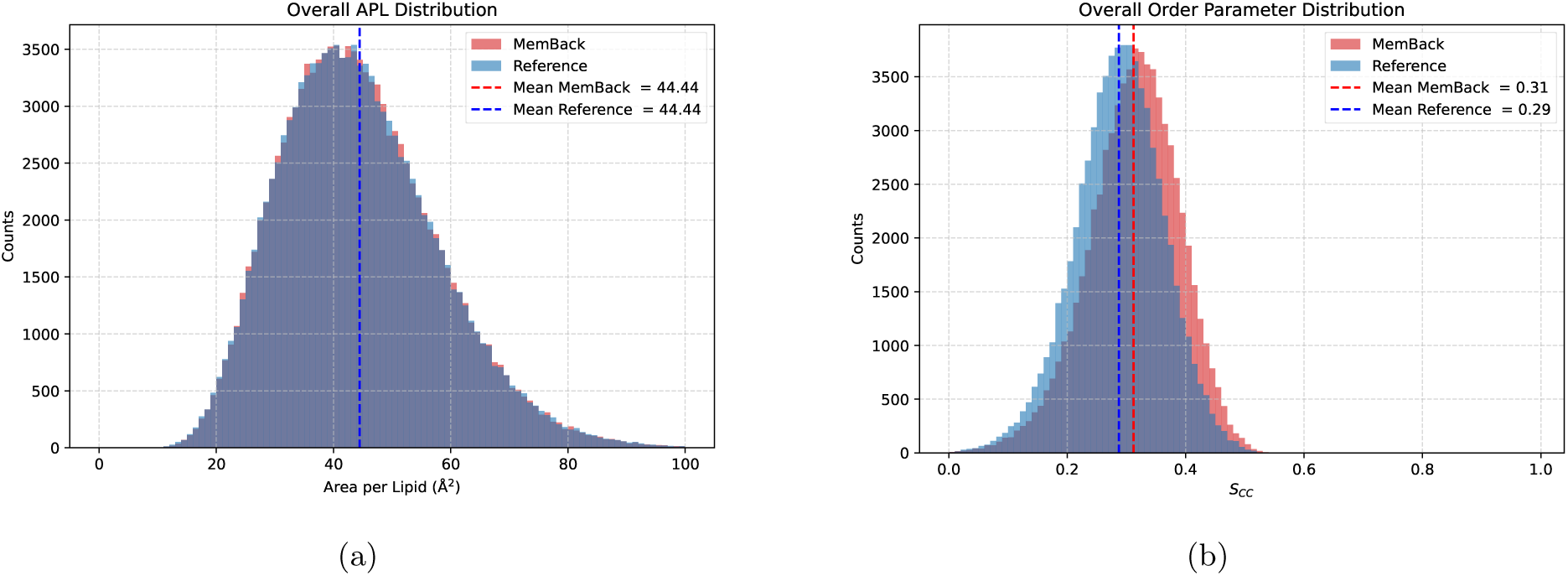
Comparison of APL and acyl chain order for 100 backmapped snapshots over 1*µs*. Backmapped results are minimized for 100 steps with position and dihedral restraints.

The order parameter distributions provide a more challenging test, since *S_CC_*depends on the reconstructed acyl-chain conformations rather than on quantities inherited from the coarse-grained input. The pooled distributions (Figure 7b) are closely superimposed, with mean values of 0.31 for MemBack and 0.29 for the reference. The reconstructed ensemble is thus slightly more ordered than the reference. A statistical comparison of MemBack and reference distributions, based on Wasserstein distances and Kolmogorov-Smirnov statistics with frame-bootstrap confidence intervals and a reference self-consistency baseline, is provided in Supporting Information Tables S3 and S4. The APL distributions differ by less than the sampling variability of the reference itself, whereas the *S_CC_* distribution is overall shifted by +0.025 across all lipid species with its width preserved.

Taken together, these results show that MemBack reconstructions reproduce the thickness, lateral packing, and chain ordering of a compositionally complex, cholesterol-rich membrane at the level of the ensemble, and that this agreement is obtained immediately after restrained minimization, without relying on extended equilibration simulations.

### 3.7 Application to Martini 3 simulations

The preceding benchmark quantifies reconstruction accuracy for forward-mapped configurations for which a corresponding atomistic reference structure is available. The intended application of MemBack, however, is the reconstruction of configurations sampled directly from production Martini 3 simulations, for which no unique atomistic reference exists. We therefore evaluated such reconstructions according to three complementary criteria: preservation of the mesoscale organization of the CG input, chemical and stereochemical integrity of the reconstructed atomistic structures, and stability during subsequent all-atom relaxation and simulation.

Three systems of increasing size and complexity were considered: a phase-separated DPPC:DLIPC:Chol bilayer containing coexisting liquid-ordered (L_o_) and liquid-disordered (L_d_) domains (Figure 8), an asymmetric red-blood-cell membrane model [27] (Figure 9), and a bicelle [31] constructed from the same multicomponent membrane composition (Figure 10). For each system, the native Martini 3 configuration was backmapped directly to CHARMM36 resolution. All three systems were subsequently energy-minimized, equilibrated, and simulated for 10 ns without restraints.

**Figure 8:**
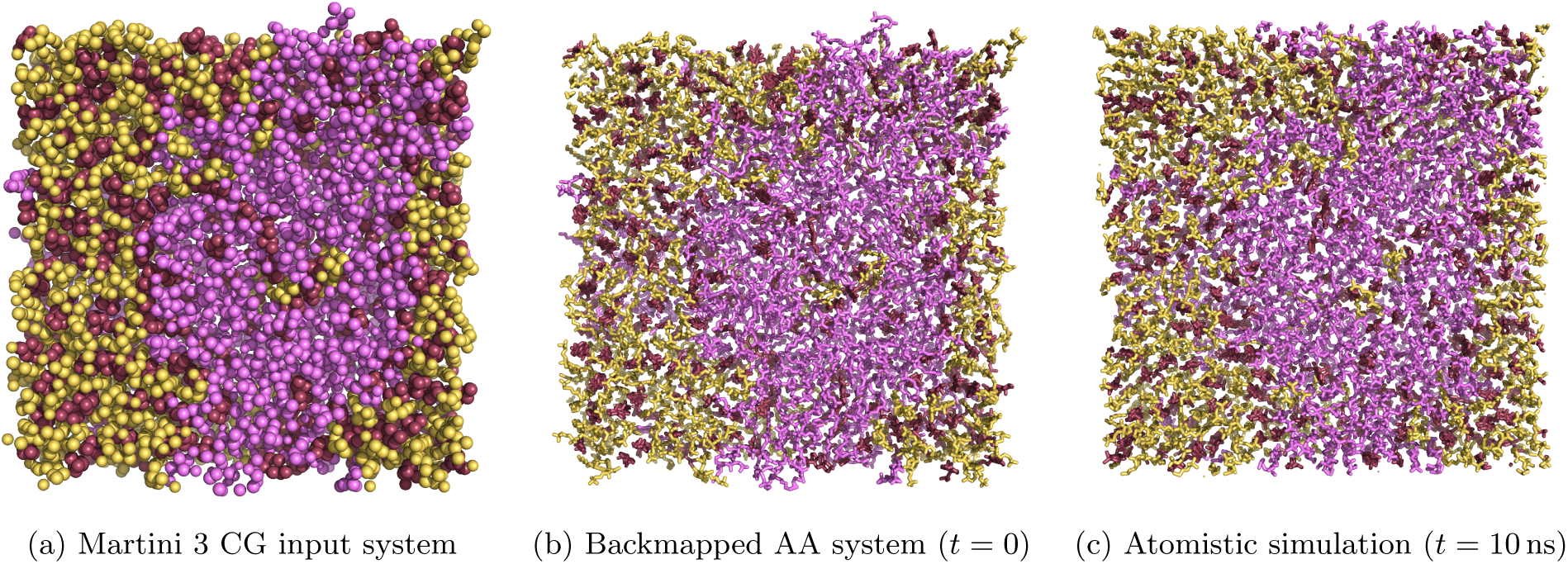
Backmapping of the phase-separated DPPC:DLIPC:CHOL membrane. DPPC, DLIPC, and cholesterol are shown in *yellow*, *magenta*, and *brown*, respectively. The L_o_/L_d_ domain organization present in the Martini 3 input (a) is retained immediately after reconstruction (b). The reconstructed system was subsequently propagated for 10 ns of unrestrained atomistic MD, confirming its stability while preserving the initial domain organization (c).

**Figure 9:**
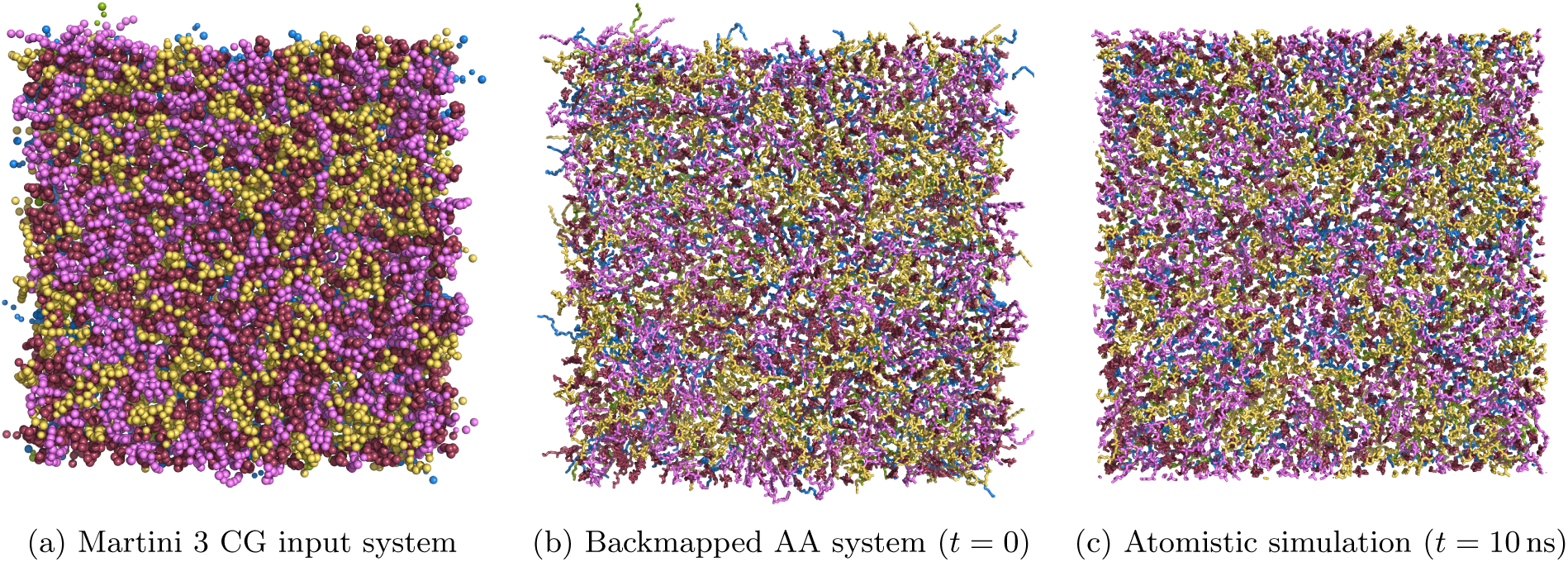
Backmapping of the asymmetric red-blood-cell membrane model. [27], viewed from the exoplasmic side. Lipid headgroups are shown according to class: PC (*yellow*), SM (*magenta*), PE (*blue*), PS (*green*), and cholesterol (*brown*). The compositional organization present in the Martini 3 configuration (a) is retained after backmapping (b). The reconstructed system was subsequently propagated for 10 ns of unrestrained atomistic MD, confirming its stability while preserving the initial domain organization (c).

**Figure 10:**
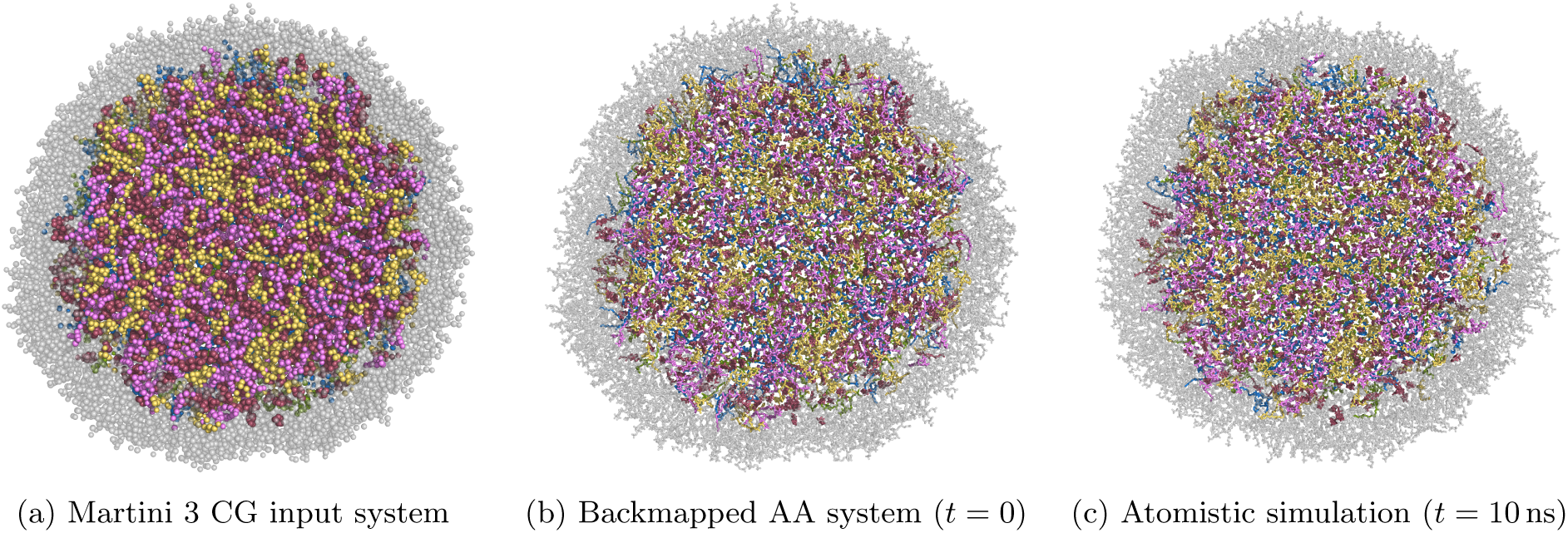
Backmapping of the multicomponent red-blood-cell bicelle. PC, SM, PE, PS, and cholesterol are shown in *yellow*, *magenta*, *blue*, *green*, and *brown*, respectively; lipids forming the outer rim are shown in semitransparent *gray*. The overall bicelle morphology and compositional organization of the Martini 3 input (a) are retained in the reconstructed atomistic system (b). The reconstructed system was subsequently propagated for 10 ns of unrestrained atomistic MD (including a flat-bottom potential, see Ref. [31] for details), confirming its stability while preserving the initial domain organization (c). bonds were classified and compared with their expected configurations. For all three systems directly upon backmapping, all chiral centers are assigned the correct handedness (100.0% over 2968 centers) and double-bond configurations were correct in 4751 of 4760 cases (99.8%). The residual is removed by the short energy minimization: after minimization both chirality and double-bond geometry are correct in 100.0% of cases. Through equilibration and production the fidelity remains high (chirality 100.0%, double bonds 100.0%). This confirms that the MemBack workflow produces chemically correct stereochemistry directly from native coarse-grained input, and that the minimization step reliably resolves the few borderline centers that arise where coarse-grained resolution underdetermines the local geometry.

Visual comparison of the CG and reconstructed configurations shows that MemBack preserves the large-scale organization encoded in the Martini 3 structures. In the DPPC:DLIPC:Chol system, the spatial separation of the L_o_- and L_d_-enriched regions is retained immediately after backmapping and remains apparent after 10 ns of atomistic simulation (Figure 8). Likewise, the compositional asymmetry and lateral organization of the red-blood-cell membrane are maintained upon reconstruction and subsequent relaxation (Figure 9). The bicelle provides a more geometrically demanding test because of its curved rim and heterogeneous lipid composition; nevertheless, its overall morphology and lipid organization are retained after reconstruction (Figure 10). Quantitative analyses of membrane structure and domain organization are presented below.

The three systems span nearly an order of magnitude in system size, from 16 541 beads for the phase-separated bilayer to 123 140 beads for the bicelle, corresponding to 182 404 and 1 417 948 atoms, respectively, after atomistic reconstruction (Table 2). Backmapping required 7.52 s, 14.25 s, and 40.13 s (including energy minimization) for the phase-separated bilayer, red-blood-cell membrane, and bicelle, respectively, on a single GPU. The reconstruction time therefore increases only moderately with system size and remains small compared with the computational cost of subsequent atomistic molecular dynamics.

**Table 2:**
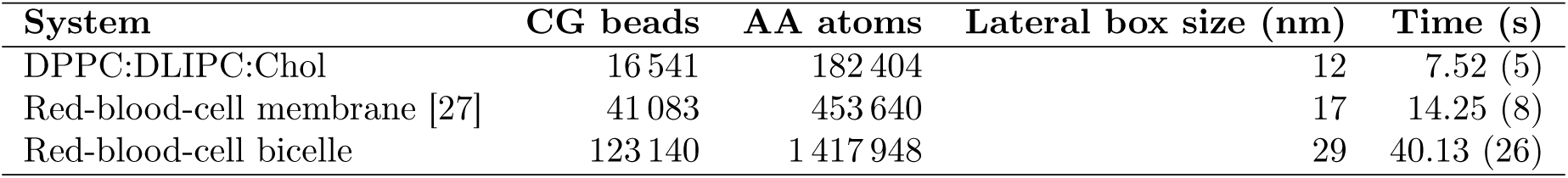
Backmapping performance for native Martini 3 test systems. The number of CG beads refers to the input Martini 3 configuration and the number of atoms to the reconstructed atomistic system. Wall-clock times report the backmapping step on a single GPU. Times in the parenthesis are times for *minimization*

| System | CG beads | AA atoms | Lateral box size (nm) | Time (s) |
| --- | --- | --- | --- | --- |
| DPPC:DLIPC:Chol | 16 541 | 182 404 | 12 | 7.52 (5) |
| Red-blood-cell membrane [27] | 41 083 | 453 640 | 17 | 14.25 (8) |
| Red-blood-cell bicelle | 123 140 | 1 417 948 | 29 | 40.13 (26) |

#### 3.7.1 Energy minimization and equilibration

Each reconstructed all-atom system was processed using the automated system-assembly, energy-minimization, and equilibration protocol described in Section 2.10. Steepest-descent minimization proceeded without numerical instability or manual intervention for all three systems. Steepest-descent minimization was run for 100 steps for the two planar bilayers and for 200 steps for the bicelle. No additional manual clash removal or modification of the reconstructed coordinates was required beyond the automated procedure described in Section 2.8.

Following minimization, the systems were subjected to the staged equilibration protocol (Supporting Information, Table S2), during which positional and stereochemical restraints were progressively released. Both planar bilayers and the bicelle subsequently entered unrestrained atomistic production simulations without instability and remained intact over the 10 ns trajectories shown in Figures 8, 9, and 10. These observations demonstrate that the reconstructed coordinates can be relaxed directly into stable atomistic simulation systems without system-specific manual intervention.

#### 3.7.2 Stereochemical fidelity on native inputs

For native Martini 3 configurations, no corresponding atomistic reference structure is available. Stereochemical fidelity was therefore assessed against the configurations specified by the chemical definition of each lipid: For every reconstructed structure, tetrahedral stereocenters and configurationally constrained double

#### 3.7.3 Round-trip consistency

As a reference-free measure of consistency with the parent CG configuration, each reconstructed structure was forward-mapped back to Martini 3 resolution and compared directly with the original CG input. For each lipid, we calculated the superposition-free RMSD between the original and reconstructed bead coordinates. A small round-trip deviation indicates that the atomistic reconstruction preserves the spatial configuration encoded by the CG representation. After energy minimization, the mean per-lipid round-trip RMSD was 0.27 Å for the DPPC:DLIPC:Chol bilayer, 0.24 Å for the asymmetric red-blood-cell membrane, and 0.28 Å for the bicelle (Figure 11). These small deviations show that the reconstructed and subsequently minimized atomistic structures remain closely consistent with their parent Martini 3 configurations. This is a necessary but not a sufficient condition for a correct reconstruction. Forward mapping averages over the atoms of each bead, so configurations differing in torsional state or stereochemistry may map onto nearly identical bead positions. The measure therefore quantifies drift away from the positional information encoded by the CG beads, and is reported only for the native Martini 3 systems, for which no atomistic reference exists. Reconstruction accuracy is instead assessed against the atomistic benchmarks of Section 3.2.

**Figure 11:**
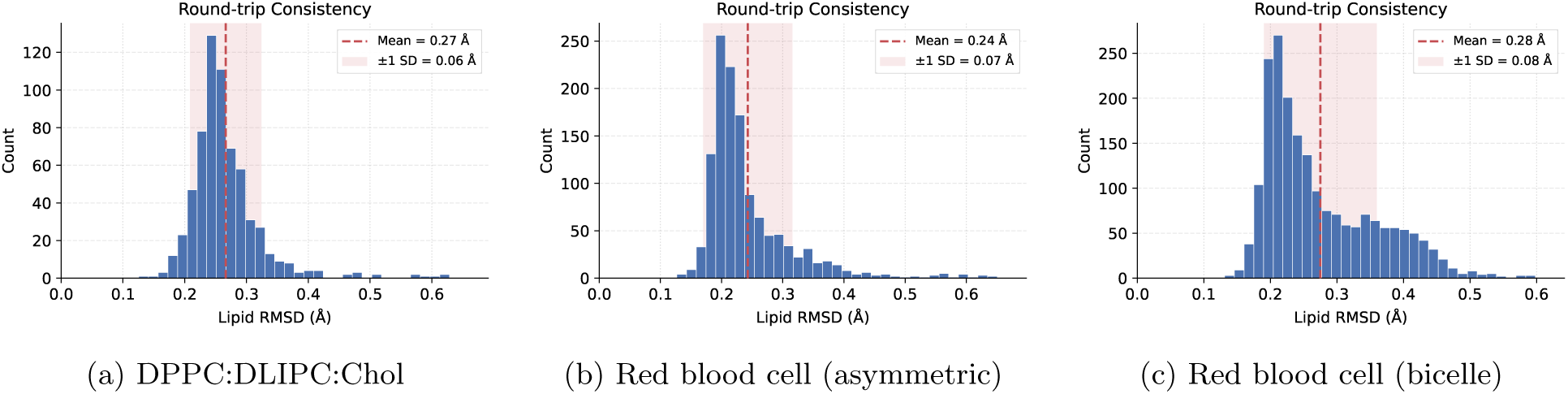
Round-trip consistency of native Martini 3 test systems after energy minimization. Each structure was backmapped to all-atom detail followed by forward-mapping back to Martini 3 resolution. The resulting bead coordinates were compared with those of the original CG input. The distributions show the superposition-free per-lipid RMSD for (a) the DPPC:DLIPC:Chol bilayer, (b) the asymmetric red-blood-cell membrane, and (c) the red-blood-cell bicelle.

#### 3.7.4 Domain analysis

The visual comparison in Figure 8 suggests that the lateral organization of the phase-separated membrane is retained upon backmapping. To quantify this preservation, we analyzed the Martini 3 and reconstructed configurations with *DomHMM* [32, 33], an unsupervised framework for identifying laterally ordered and disordered membrane domains. *DomHMM* assigns each lipid an ordered or disordered state using a Gaussian hidden Markov model trained per lipid type on the area per lipid and the average acyl-chain order parameter, and subsequently groups lipids into contiguous lateral domains by spatial autocorrelation of these states followed by hierarchical clustering. Because the classification relies only on lipid coordinates, the same analysis can be applied to coarse-grained and atomistic trajectories, which makes it suitable for a direct comparison across resolutions.

One hundred configurations were extracted from the Martini 3 DPPC:DLIPC:Chol trajectory between 9 and 10 µs at intervals of 10 ns. Each configuration was independently backmapped with MemBack and energy-minimized as described in Section 2.10; no subsequent equilibration or production runs were performed. *DomHMM* was applied independently to the Martini 3 and corresponding backmapped configurations, with the two membrane leaflets analyzed separately. Because lipid identities are retained during backmapping, domain assignments can be compared on a lipid-by-lipid basis between the two resolutions. For each frame and leaflet, agreement between the coarse-grained and backmapped assignments was quantified using the Pearson correlation coefficient, resulting in 200 comparisons for each domain class.

The resulting domain maps show close agreement in their location, shape, and lateral extent (Figure 12). Across all frames and leaflets, the correlation between the coarse-grained and backmapped domain assignments was 0.89*±*0.08 for ordered domains and 0.90*±*0.06 for disordered domains (mean *±* standard deviation; Figure 13). The distributions are strongly concentrated at high correlation values, with most comparisons exceeding 0.8.

**Figure 12:**
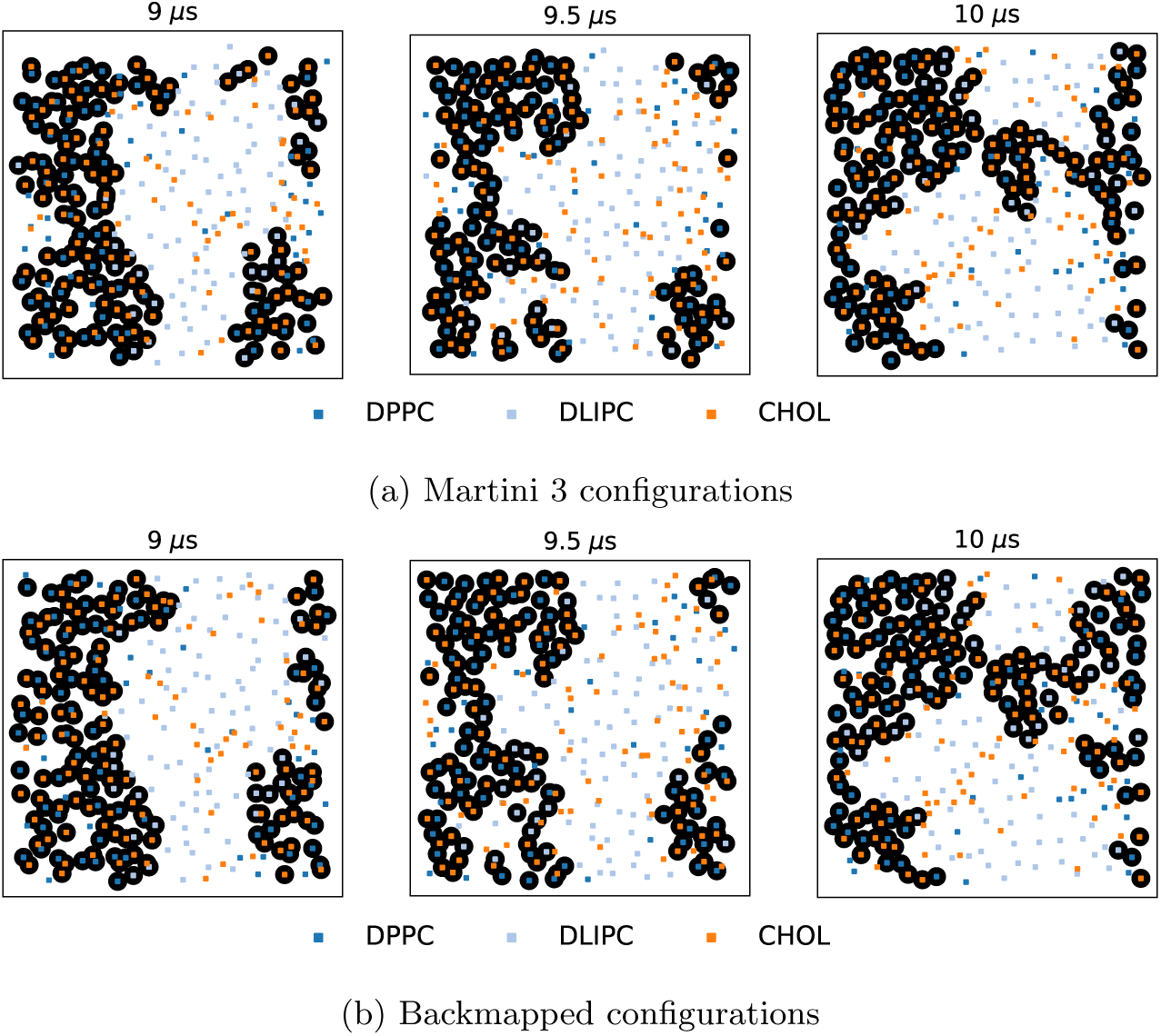
Preservation of ordered membrane domains upon backmapping. Ordered domains identified by *DomHMM* are shown for three representative configurations of the DPPC:DLIPC:Chol membrane. Small markers denote individual lipids colored by species, whereas circled markers indicate lipids assigned to an ordered domain. The location, shape, and extent of the domains identified in the Martini 3 configurations (a) are closely reproduced after backmapping and energy minimization (b), without subsequent atomistic equilibration.

**Figure 13:**
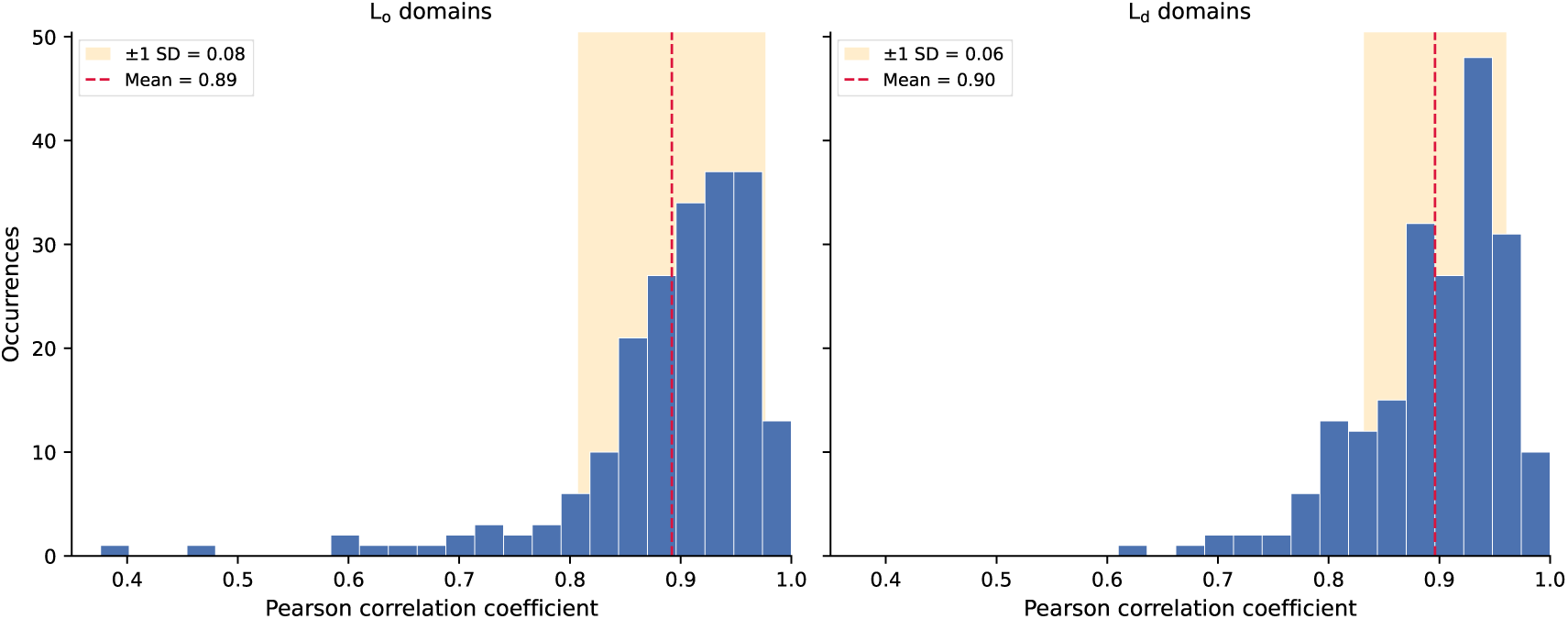
Agreement between domain assignments before and after backmapping. Distributions of the Pearson correlation coefficient between lipid-wise domain assignments in the Martini 3 DPPC:DLIPC:Chol configurations and their corresponding backmapped and energy-minimized structures. Correlations are shown separately for ordered (*left*) and disordered (*right*) domains and comprise 100 configurations analyzed independently for both leaflets (*N* = 200 per domain class).

The remaining discrepancies occur predominantly for lipids close to domain boundaries, where small changes in local packing or chain conformation can alter the domain classification. This sensitivity is expected because the *DomHMM* assignment depends on area per lipid and acyl-chain ordering, neither of which is imposed directly during reconstruction. Rather, both emerge from the reconstructed atomistic coordinates. The high agreement therefore shows that MemBack preserves not only the positions of individual lipids but also the mesoscale lateral organization encoded in the parent Martini 3 configuration. Importantly, this correspondence is observed directly after backmapping and restrained energy minimization, without atomistic equilibration that could otherwise reorganize or restore the domain structure.

### 3.8 Discussion and conclusions

In this work, we introduced *MemBack*, an SE(3)-equivariant graph neural network for reconstructing CHARMM36 lipid systems from Martini 3 coarse-grained configurations. Heavy-atom coordinates are generated in a single network evaluation as bead-relative displacement vectors, followed by automated hydrogen reconstruction, solvent and ion conversion, stereochemical correction, clash resolution, and system assembly. Across chemically diverse forward-mapped configurations, *MemBack* achieved a mean superposition-free per-lipid heavy-atom RMSD of 0.65 Å, compared with 1.04 Å for CG2AT2, while reproducing bond lengths, bond angles, and stereochemically relevant dihedrals with mean errors of 0.03 Å, 4.3*^◦^*, and 6.5*^◦^*, respectively.

Comparable accuracy was retained for membrane systems and lipid species excluded from training, showing that performance is not restricted to interpolation within the training systems.

For multiscale simulation, however, reconstruction quality is not defined solely by agreement with a particular atomistic reference, since a native CG configuration is compatible with multiple atomistic realizations. More relevant is whether the reconstructed system preserves the physical state sampled at coarse-grained resolution. In an RBC membrane excluded from training, bilayer thickness, area per lipid, and acyl-chain order closely matched the atomistic reference after only restrained minimization. Native Martini 3 configurations likewise remained close to their parent CG structures, with round-trip deviations of only 0.24–0.28 Å. In the phase-separated DPPC:DLIPC:Chol membrane, ordered and disordered domains were also preserved after reconstruction, with mean lipid-wise assignment correlations of 0.89 *±* 0.08 and 0.90 *±* 0.06, respectively.

Because these comparisons were performed before atomistic equilibration, the agreement reflects information retained during backmapping rather than subsequent recovery by CHARMM36 dynamics.

The successful reconstruction of DLIPC, PLA20, and PDOPE further indicates that *MemBack* learns transferable local structural environments rather than complete lipid identities. Lipid residue identity is not supplied to the network; reconstruction is conditioned instead on Martini bead chemistry, bonded connectivity, and local molecular geometry. The approximately 2.7 million lipid graphs from 50 species and about 58 µs of atomistic simulation therefore provide a basis for combining recurring headgroup, backbone, and acyl-chain motifs in previously unseen lipids. This transfer remains limited to bead types and local bonded motifs represented during training and does not establish unrestricted extrapolation to novel chemical environments.

Compared with fragment-based approaches, *MemBack* does not require a curated library of molecular conformers or iterative fragment fitting during inference. This distinction is relevant to the comparison with CG2AT2, which used fragment definitions derived from atomistic simulations closely related to the benchmark systems and thus a favorable fragment basis. Nevertheless, *MemBack* yielded lower reconstruction errors across all systems. With default CG2AT2 fragments, errors increased further for several lipids, illustrating the dependence of fragment-based reconstruction on representative conformers and the associated system-specific preparation. At the same time, the comparison does not constitute an exhaustive evaluation of all possible CG2AT2 parameterizations. Reconstruction nevertheless still requires a mapping that assigns the atoms of each lipid species to its coarse-grained beads, which defines the identity of the atoms to be placed and is therefore needed for every species, including those absent from training. To keep this requirement from limiting applicability, *MemBack* is distributed with a mapping database covering 150 lipid species and with a tool that derives the mapping of a new lipid directly from its Martini 3 and CHARMM36 topologies.

The local molecular representation also makes *MemBack* naturally scalable: each lipid is reconstructed independently, so graph size is determined by the molecule rather than the complete membrane. This enabled reconstruction of systems ranging from planar bilayers to a multicomponent bicelle containing more than 1.4 million atoms while preserving their overall morphology and compositional organization. The same locality, however, means that the network does not explicitly account for neighboring molecules and cannot enforce intermolecular steric compatibility during inference. Residual overlaps are therefore removed during automated post-processing and restrained minimization. Incorporating short-range intermolecular information could address this limitation, although at the cost of increased graph size and reduced molecule-wise scalability.

A further limitation follows from the intrinsically many-to-one nature of coarse-graining. *MemBack* generates a single atomistic reconstruction for each CG input, whereas multiple atomistic conformations may be compatible with the same Martini configuration. Deterministic regression may therefore reduce part of the conformational variability lost during coarse-graining; the slightly increased acyl-chain order in the reconstructed RBC membrane is consistent with, but does not establish, such an effect. Conditional generative approaches could instead sample multiple atomistic realizations. The present implementation is additionally restricted to lipid systems and to the Martini 3-to-CHARMM36 transformation; application to different molecular classes, CG mappings, or atomistic force fields would require retraining or adaptation.

Taken together, the results establish *MemBack* as an accurate and scalable interface between Martini 3 and CHARMM36 membrane simulations. It combines superposition-free sub-Å reconstruction accuracy with transfer to unseen membrane compositions and lipid species, preservation of membrane-scale organization with minimal relaxation, and reconstruction of systems containing more than one million atoms. These properties enable configurational states sampled at coarse-grained resolution to be transferred efficiently to atomistic resolution without extensive system-specific preparation.

## Data Availability Statement

Figures and tables that could not be included in the body are provided in the Supporting Information. *MemBack* is made available on the research group’s GitHub repository (https://github.com/BioMemPhys-FAU/memback).

## Supporting Information

Convergence of the production model during training; Accuracy of hydrogen atom reconstruction; Set of membrane systems and simulations employed in training of the MemBack model; six-stage protocol adapted from the standard CHARMM-GUI [22];

## Acknowledgements

We thank Cristian Popov, Nico Piontek, and Marius Trollmann for providing simulation data for several membrane systems used in this study. This work was supported by the German Research Foundation (Deutsche Forschungsgemeinschaft, DFG) through grants 534019561 and 555589280. We also gratefully acknowledge the scientific support and HPC resources provided by the Erlangen National High Performance Computing Center (NHR@FAU) of the Friedrich-Alexander-Universität Erlangen-Nürnberg (FAU). The hardware is partially funded by the German Research Foundation (DFG).

---

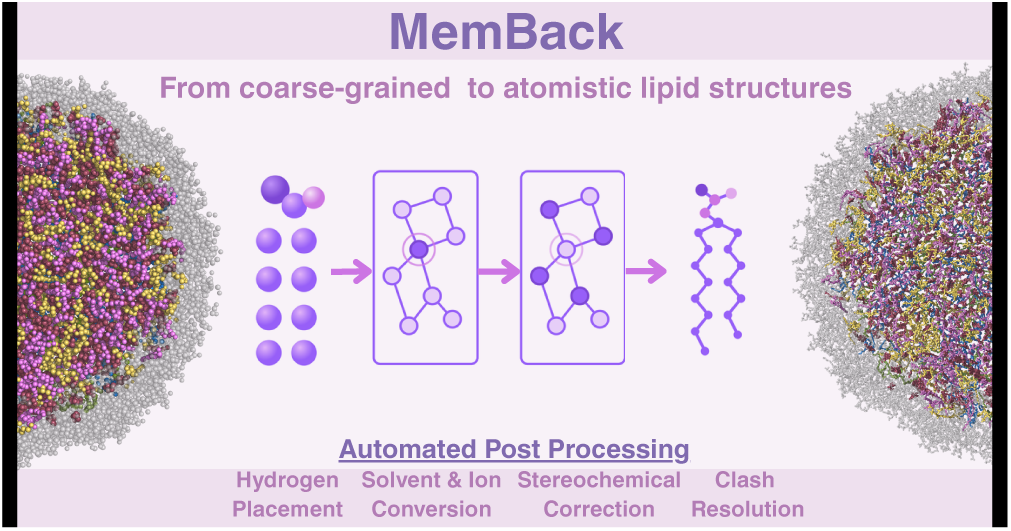

