## Supporting Information for "MemBack: An Equivariant Graph Neural Network for Backmapping Lipid Membranes"

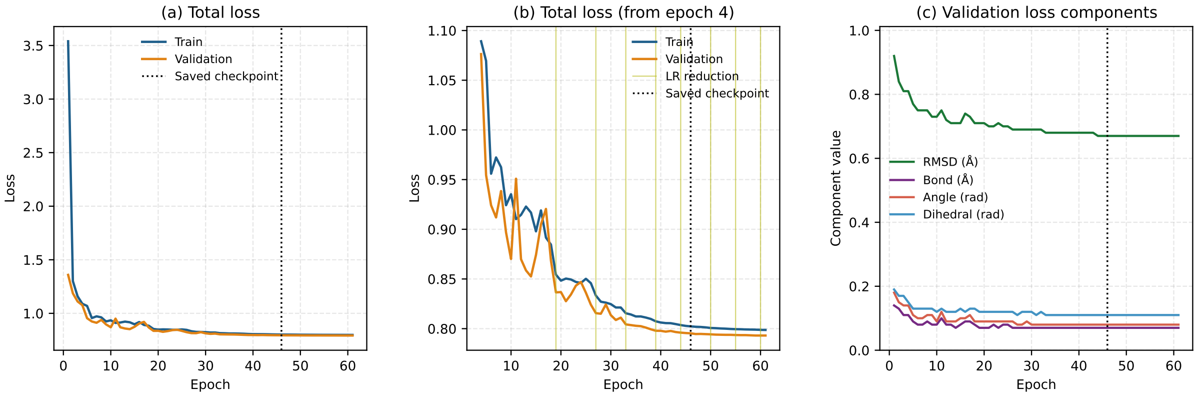


Figure S1: **Convergence of the production model during training.** (a) Training and validation loss over the complete optimization trajectory. (b) Expanded view from epoch 4 onward. Vertical lines indicate learning-rate reductions, and the dotted line marks the retained checkpoint at epoch 46. (c) Validation metrics for the coordinate, bond, angle, and torsional contributions to the objective. The coordinate RMSD decreases rapidly during the initial optimization and subsequently approaches a plateau, while the local-geometry terms converge concurrently. Training and validation losses remain closely matched throughout optimization.

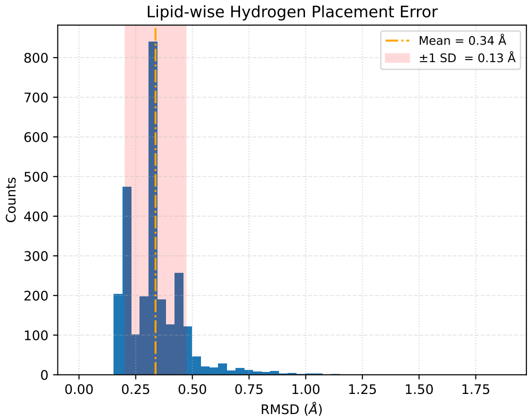


Figure S2: **Accuracy of hydrogen atom reconstruction.** The distribution shows the per-lipid RMSD between reconstructed and reference hydrogen atom coordinates, pooled over one frame each from PSM:POPC:Chol, DMPA:DMPE, DPPC, and the multicomponent red-blood-cell membrane. Heavy-atom coordinates were kept fixed at their reference positions to isolate the hydrogen-placement step.

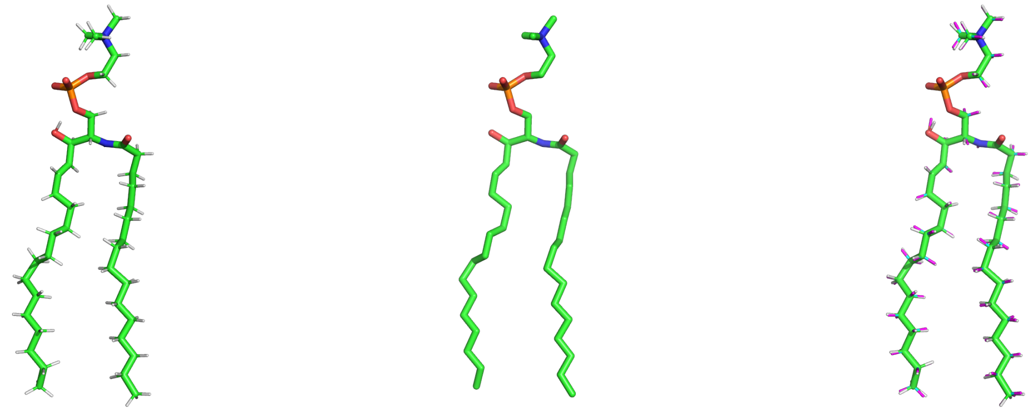

(a) (b) (c)

Figure S3: **Reconstruction of hydrogen positions.**(a) Reference all-atom configuration of a PSM molecule taken from a CHARMM36 trajectory.(b) The same configuration after removal of all hydrogen atoms, which constitutes the input to the hydrogen-placement procedure. (c) Superposition of the reference hydrogen positions (*white*) and those reconstructed from the unchanged heavy-atom coordinates (*magenta*). The heavy-atom coordinates are identical in all three panels. Because the reference hydrogens are configurations sampled during molecular dynamics whereas the reconstruction places them at idealized bond lengths and angles, the two sets do not coincide exactly.

Table S1: Composition of the simulation database used to construct the MemBack training set. Counts are given per lipid/water/ion species in the order listed under Composition; box sizes are lateral (x-y) dimensions. Blank entries indicate metadata not tabulated in the source records. For the red blood cell membrane composition only the total number of lipids is given (12 different lipid types). All simulations are neutralized with 0.15 m NaCl.

| **System** | **Composition** | **# of Atoms** | **Box(x-y)** | **Time** | **Temp.** |
|---|---|---|---|---|---|
| PSM:POPC:Chol | 468:468:234 | 139 464 | 17.5 nm | 2.7 μs | 303.15 K |
| DPPC | 100 | 13 000 | 5.5 nm | 500 ns | 325 K |
| DMPC | 200 | 23 600 | 7.9 nm | 500 ns | 325 K |
| DMPE | 200 | 21 800 | 7.2 nm | 500 ns | 325 K |
| Red Blood Cell [1] | 1256 | 116 808 | 17.5 nm | 20 μs | 310 K |
| DOPA:DOPE | 88:88 | 21 912 | 7.5 nm | 1.0 μs | 300 K |
| DOPC:DOPE | 84:84 | 22 428 | 7.5 nm | 1.0 μs | 300 K |
| DOPE:DOPS | 84:84 | 21 840 | 7.5 nm | 1.0 μs | 300 K |
| DOPG:DOPE | 86:86 | 22 360 | 7.5 nm | 1.0 μs | 300 K |
| PAPA:POPE | 48:142 | 23 318 | 7.5 nm | 1.0 μs | 300 K |
| PAPE:POPE | 48:142 | 23 750 | 7.5 nm | 1.0 μs | 300 K |
| PLPG:PLPE | 46:136 | 22 478 | 7.5 nm | 1.0 μs | 300 K |
| POPA | 188 | 21 808 | 7.5 nm | 1.0 μs | 300 K |
| POPE:POPS | 94:94 | 23 688 | 7.5 nm | 1.0 μs | 300 K |
| POPG:POPE | 138:46 | 23 276 | 7.5 nm | 1.0 μs | 300 K |
| POPI:POPE | 38:148 | 23 706 | 7.5 nm | 1.0 μs | 300 K |
| SAPC:POPE:Chol | 66:66:66 | 22 374 | 7.5 nm | 1.0 μs | 300 K |
| SAPE:SAPS:Chol | 66:66:66 | 22 308 | 7.5 nm | 1.0 μs | 300 K |
| SAPI:POPE | 38:148 | 23 934 | 7.5 nm | 1.0 μs | 300 K |
| SAPI:POPE:Chol | 50:100:50 | 23 350 | 7.5 nm | 1.0 μs | 300 K |
| SOPE:POPS | 96:96 | 24 768 | 7.5 nm | 1.0 μs | 300 K |
| PLPI:POPE | 38:148 | 23 630 | 7.5 nm | 1.0 μs | 300 K |
| SDPE:SDPS | 88:88 | 23 584 | 7.5 nm | 1.0 μs | 300 K |
| SOPG | 166 | 22 078 | 7.5 nm | 1.0 μs | 300 K |
| TOCL2:POPE:POPC | 16:64:80 | 22 688 | 7.5 nm | 1.0 μs | 300 K |
| SDPA:SDPE | 44:132 | 23 012 | 7.5 nm | 1.0 μs | 300 K |
| SDPI:POPE:Chol | 40:118:40 | 23 510 | 7.5 nm | 1.0 μs | 300 K |
| SOPA | 192 | 23 424 | 7.5 nm | 1.0 μs | 300 K |
| POPI33:POPE:POPS:Chol | 18:102:34:52 | 23 562 | 7.5 nm | 1.0 μs | 310 K |
| POPI2C:POPE:POPS | 18:112:56 | 23 704 | 7.5 nm | 1.0 μs | 310 K |
| POPI15:POPE | 18:170 | 23 770 | 7.5 nm | 1.0 μs | 310 K |
| SAPI13:POPE:POPS | 18:112:56 | 23 740 | 7.5 nm | 1.0 μs | 310 K |
| SAPI2A:POPE | 18:170 | 23 950 | 7.5 nm | 1.0 μs | 310 K |
| SAPI33:POPE:POPS:Chol | 18:102:34:52 | 23 670 | 7.5 nm | 1.0 μs | 310 K |
| DMPA:DMPE | 92:92 | 19 228 | 7.5 nm | 1.0 μs | 320 K |
| DMPG:DMPE:Chol | 70:70:70 | 20 580 | 7.5 nm | 1.0 μs | 320 K |
| TMCL2:DPPE:DPPC | 20:114:38 | 22 894 | 7.5 nm | 1.0 μs | 350 K |

Table S2: Six-stage equilibration protocol. Position restraints act on the headgroup atoms of the lipids (phosphorus atom for phospholipids, oxygen atom for cholesterol) and dihedral restraints on the stereochemistry-determining centers; both are released progressively. Pressure coupling (C-rescale [2], semi-isotropic, 1 bar) is active from stage 3.

| Stage | Ensemble | $\Delta t$ (fs) | Length (ns) | POSRES (kJ mol$^{-1}$ nm$^{-2}$) | DIHRES (kJ mol$^{-1}$ rad$^{-2}$) |
|---|---|---|---|---|---|
| 1 | NVT | 1 | 0.125 | 1000 | 1000 |
| 2 | NVT | 1 | 0.125 | 400 | 400 |
| 3 | NPT | 1 | 0.125 | 400 | 200 |
| 4 | NPT | 2 | 0.500 | 200 | 200 |
| 5 | NPT | 2 | 0.500 | 40 | 100 |
| 6 | NPT | 2 | 0.500 | 0 | 0 |

Table S3: **Statistical comparison of the pooled area per lipid (APL) and acyl-chain order parameter ($S_{CC}$) distributions.** Values are pooled over all lipids and 100 backmapped snapshots of the red-blood-cell membrane ($n$ = 116,800 for APL and $n$ = 69,200 for $S_{CC}$ per data set). $W_1$ is the Wasserstein-1 distance between the MemBack and reference distributions, in the units of the observable. $D$ is the two-sample Kolmogorov–Smirnov statistic, i.e. the maximum difference between the cumulative distributions. 95% confidence intervals were obtained by bootstrap resampling of frames (500 resamples), which accounts for correlations between lipids within a frame. Because the pooled samples comprise $\sim 10^5$ correlated observations, significance tests reject the null hypothesis for arbitrarily small deviations; agreement is therefore assessed with these effect sizes relative to a self-consistency baseline. The baseline columns compare the even-numbered reference frames with the odd-numbered reference frames (reference sampling variability) and the even-numbered MemBack frames with the odd-numbered reference frames, at identical sample sizes. The mean APL is fixed by the box dimensions inherited from the coarse-grained input and is therefore identical by construction. For $S_{CC}$, the per-frame mean difference is $+0.0249 \pm 0.0013$ and the per-frame difference in standard deviation is $+0.0021 \pm 0.0012$ (mean ± SD over 100 frames), indicating a uniform shift of the distribution without a change in its width.

| | Mean / SD | | MemBack vs. reference (all frames) | | Baseline $W_1$ (even vs. odd frames) | |
|---|---|---|---|---|---|---|
| Observable | MemBack | Reference | $W_1$ [95% CI] | $D$ [95% CI] | Ref. vs. Ref. | MemBack vs. Ref. |
| APL (Å$^2$) | 44.44 / 13.47 | 44.44 / 13.54 | 0.043 [0.020, 0.127] | 0.002 [0.001, 0.004] | 0.142 | 0.101 |
| $S_{CC}$ | 0.312 / 0.077 | 0.287 / 0.075 | 0.0249 [0.0246, 0.0252] | 0.148 [0.146, 0.151] | 0.0004 | 0.0252 |

Table S4: **Per-lipid-species comparison of APL and $S_{CC}$ distributions** for the 100 backmapped snapshots of the red-blood-cell membrane. $W_1$, Wasserstein-1 distance; $D$, two-sample Kolmogorov–Smirnov statistic; $\Delta\mu$, difference of means (MemBack - reference). Unlike the overall mean APL, the partitioning of area among lipid species is not constrained by the box dimensions. $S_{CC}$ is the average over both acyl chains (for sphingolipids, the sphingosine and amide-linked chains) and is not defined for cholesterol. *, lipid species absent from the training database.

| | APL (Å$^2$) | | | $S_{CC}$ | | |
|---|---|---|---|---|---|---|
| Lipid | $W_1$ | $D$ | $\Delta\mu$ | $W_1$ | $D$ | $\Delta\mu$ |
| DPPC | 0.17 | 0.015 | +0.06 | 0.023 | 0.168 | +0.023 |
| PAPC | 0.16 | 0.011 | −0.13 | 0.028 | 0.200 | +0.028 |
| PLPC | 0.10 | 0.005 | −0.03 | 0.026 | 0.174 | +0.026 |
| POPC | 0.14 | 0.011 | +0.08 | 0.031 | 0.211 | +0.031 |
| SOPC | 0.19 | 0.009 | −0.14 | 0.031 | 0.208 | +0.031 |
| PDOPE* | 0.23 | 0.015 | +0.20 | 0.024 | 0.169 | +0.024 |
| POPE | 0.23 | 0.016 | +0.19 | 0.028 | 0.174 | +0.028 |
| SAPE | 0.16 | 0.009 | +0.12 | 0.027 | 0.180 | +0.026 |
| PLA20* | 0.15 | 0.007 | +0.13 | 0.024 | 0.168 | +0.024 |
| PAPS | 0.30 | 0.013 | +0.29 | 0.027 | 0.183 | +0.027 |
| SAPS | 0.22 | 0.016 | +0.17 | 0.028 | 0.201 | +0.028 |
| SAPI2A | 0.66 | 0.028 | +0.66 | 0.026 | 0.188 | +0.026 |
| LSM | 0.16 | 0.009 | −0.04 | 0.016 | 0.102 | +0.016 |
| NSM | 0.22 | 0.011 | −0.17 | 0.018 | 0.128 | +0.018 |
| PSM | 0.11 | 0.006 | −0.05 | 0.025 | 0.171 | +0.025 |
| CHL1 | 0.08 | 0.005 | −0.06 | – | – | – |

| | **SCD model**<br>symmetric cholesterol distribution | **ACD model**<br>asymmetric cholesterol distribution | |
|---|---|---|---|
| *System description* | | | |
| Lipid species | 16 | 12 | |
| Total lipids | 1168 | 1256 | |
| *Transbilayer organization (exoplasmic / cytoplasmic)* | | | |
| Cholesterol[a] | 476 (240 / 236) | 520 (401 / 119) | |
| Phospholipids | 692 (360 / 332) | 736 (320 / 416) | |
| All lipids | 1168 (600 / 568) | 1256 (721 / 535) | |
| Cholesterol per leaflet (mol%) | 40 / 42 | 56 / 22 | |
| Phospholipid imbalance | +8% (exoplasmic) | +30% (cytoplasmic) | |
| *Lipid composition, total (exoplasmic / cytoplasmic)* | | | In training database |
| CHL1 | 476 (240/236) | 520 (401/119) | yes[b] |
| DPPC | 12 (0/12) | - | yes |
| LSM | 48 (48/0) | 48 (48/0) | yes |
| NSM | 56 (56/0) | 56 (56/0) | yes |
| OAPE | - | 32 (0/32) | yes |
| PAPC | 32 (32/0) | 24 (24/0) | yes |
| PAPS | 76 (0/76) | 120 (0/120) | yes |
| PDOPE | 44 (0/44) | 72 (0/72) | **no**[c] |
| PLA20 | 84 (16/68) | - | **no** |
| PLPC | 140 (88/52) | 160 (80/80) | yes |
| PLQS | - | 80 (0/80) | **no**[c] |
| POPC | 20 (0/20) | 32 (0/32) | yes |
| POPE | 12 (0/12) | - | yes |
| PSM | 80 (72/8) | 72 (72/0) | yes |
| SAPE | 20 (0/20) | - | yes |
| SAPI2A | 12 (0/12) | - | yes |
| SAPS | 16 (8/8) | - | yes |
| SOPC | 40 (40/0) | 40 (40/0) | yes |

Table S5: **Comparison of the two red-blood-cell membrane models used in this work.** Both trajectories were taken from ref. [3]; the SCD model is based on the lipidomics data of Lorent *et al.* [4] and the ACD model on the leaflet compositions reported in Doktorova *et al.* [1]. The two models describe the same human erythrocyte lipidome but differ in their transbilayer organization. The SCD model has comparable phospholipid and cholesterol numbers in the two leaflets, whereas the ACD model combines a 30% phospholipid excess in the cytoplasmic leaflet with a strong cholesterol overabundance within the exoplasmic leaflet. Leaflet assignment was determined from the phosphate atoms of the phospholipids and the hydroxyl oxygen of cholesterol. The last column indicates whether atomistic simulation data for the respective lipid type entered the training database. The SCD membrane was excluded from training in its entirety. [a]Cholesterol counts are for the analyzed configuration because cholesterol translocates between leaflets, the instantaneous split varies. [b]Cholesterol was excluded when extracting training data from the ACD membrane. Its training data originate from the other bilayers listed in Table S1. [c]Present in the ACD membrane but excluded when extracting training data from it.
